# The effects of polydisperse crowders on the compaction of the *Escherichia coli* nucleoid

**DOI:** 10.1101/803130

**Authors:** Da Yang, Jaana Männik, Scott T. Retterer, Jaan Männik

## Abstract

DNA binding proteins, supercoiling, macromolecular crowders, and transient DNA attachments to the cell membrane have all been implicated in the organization of the bacterial chromosome. However, it is unclear what role these factors play in compacting the bacterial DNA into a distinct organelle-like entity, the nucleoid. By analyzing the effects of osmotic shock and mechanical squeezing on *Escherichia coli*, we show that macromolecular crowders play a dominant role in the compaction of the DNA into the nucleoid. We find that a 30% increase in the crowder concentration from physiological levels leads to a 3-fold decrease in the nucleoid’s volume. The compaction is anisotropic, being higher along the long axes of the cell at low crowding levels. At higher crowding levels the compression becomes isotropic, implying that *E. coli* nucleoids lack a well-defined backbone. We furthermore show that the compressibility of the nucleoid is not significantly affected by cell growth rates and by prior treatment with rifampicin. The latter results point out that in addition to poly-ribosomes, soluble cytoplasmic proteins have a significant contribution in determining the size of the nucleoid.

## INTRODUCTION

Fully replicated chromosomal DNA in *Escherichia coli* consists of approximately 4.6 Mbp of DNA and has a contour length of 1.6 mm. Despite being about a thousand times longer than the dimensions of the cell, it is confined to a fraction of the cytosolic volume, which is referred to as the nucleoid (Wang *et al.*, 2013). The cytosolic volume fraction occupied by a nucleoid has been estimated to be in the range of 20%-75% in *E. coli* (Woldringh, 2002, Sherratt, 2003, Fisher *et al.*, 2013) and many other bacterial species (Gray *et al.*, 2019). In contrast, when DNA is released from the cell, it expands to a volume that exceeds the cell volume by about 1000-fold (Cunha *et al.*, 2001, Romantsov *et al.*, 2007, Wegner *et al.*, 2012). What leads to the significant compaction of the DNA within the cell is not completely understood. Answering this question is important because DNA compaction can be expected to affect DNA replication, segregation, repair, and transcription. Via transcription, the size of the nucleoid can exert control over a majority of cellular activities.

Several factors contributing to the size of the nucleoid have been proposed. These include nucleoid associated proteins (NAPs), DNA supercoiling, DNA linkages to the membrane (transertion), transcriptional activity, and molecular crowders (Wang *et al.*, 2013, de Vries, 2010, Jin *et al.*, 2013). How much each of these factors/processes contributes to compacting the millimeter-long DNA into the micron-sized cell has remained unclear. Although several NAPs, which can bridge (H-NS) and bend (HU, Fis, IHF) chromosomal DNA, are abundantly present in *E. coli* during its lag-phase growth (for review see (Dame *et al.*, 2011, Dillon and Dorman, 2010)) they appear to affect chromosome conformations only at the local scale, in regions spanning less than 300 kb (Lioy *et al.*, 2018). Moreover, the removal of these binding proteins from the cell one at a time does not change the nucleoid size (Wu *et al.*, 2019, Wang *et al.*, 2013). The exception is a low-abundance of MatP that anchors the replication terminus region to the divisome (Espeli *et al.*, 2012, Männik *et al.*, 2016) and conveys different organization to this ~800 kb long domain (Mercier *et al.*, 2008, Lioy *et al.*, 2018). Along with NAPs, DNA supercoiling can also be expected to compact chromosomal DNA (Woldringh *et al.*, 1995). It has been proposed that individual supercoils can nematically align at densities, 13 g/l, which could lead to a compacted polymer (Odijk, 1998). Although such DNA densities are present in the *E. coli* nucleoid, correlations between DNA supercoiling levels and nucleoid size have not been directly observed yet (Stuger *et al.*, 2002, Romantsov *et al.*, 2007, Cagliero and Jin, 2013). However, it has been reported that transcription leads to significant compaction of the nucleoid in fast growth conditions (Cagliero and Jin, 2013). This compaction could be explained by an increased level of supercoiling due to transcription, but alternative mechanisms cannot be ruled out yet.

In addition to the mechanisms that compact the nucleoid, an active process, termed transertion, expands it (Libby *et al.*, 2012, Bakshi *et al.*, 2014, Bakshi *et al.*, 2012). Transertion refers to a process whereby DNA is tethered to the plasma membrane via concurrently occurring transcription, translation, and membrane insertion (Woldringh, 2002, Roggiani and Goulian, 2015). Transertion linkages can form when an integral membrane protein with an N-terminal membrane domain is synthesized. These linkages consist of RNA polymerase, transcribed mRNA, translating ribosomes, and the membrane protein that has inserted its N-terminal domain into the cell membrane before its synthesis has completed. These large and structurally complex linkages are inherently short-lived and limited by the time it takes to transcribe a gene (~10 sec). Evidence for the existence of these linkages has been provided in recent studies (Libby *et al.*, 2012, Bakshi *et al.*, 2014), but the conditions in which the linkages affect nucleoid size have not been mapped out yet.

While all the above processes have been extensively studied, none of them appear to be capable of explaining the extent of the compaction needed to confine DNA into the experimentally observed nucleoid sizes. A number of theoretical (Odijk, 1998) and modeling studies (Mondal *et al.*, 2011, Shendruk *et al.*, 2015, Shin *et al.*, 2014, Kim *et al.*, 2015, Joyeux, 2018) have pointed out the importance of macromolecular crowders on the compaction of chromosomal DNA. In the seminal work, Odijk proposed that crowders and DNA separate into two distinct phases in an *E. coli* cell, a nucleoid and cytosolic phase (Odijk, 1998). The nucleoid phase contains the chromosome and is depleted of cytoplasmic proteins, whereas the cytosolic phase has an excess number of soluble proteins. Several modeling studies have confirmed these predictions (Mondal *et al.*, 2011, Shendruk *et al.*, 2015, Shin *et al.*, 2014, Kim *et al.*, 2015). However, all these results are based on equilibrium thermodynamics and coarse-grained models of the DNA and the cytosol. Therefore, the validity of underlying assumptions can be hardly taken for granted.

To date, quantitative experimental tests to verify these models have been carried out *in vitro*, using purified DNA (Zhang *et al.*, 2009) or DNA liberated from cells (Cunha *et al.*, 2001, Pelletier *et al.*, 2012). In these studies, charge-neutral polymers, dextran or polyethylene glycol (PEG), have been used as crowding agents to mimic the cytoplasmic environment even though most cytosolic crowders do not have a neutral charge. All these experiments agree that crowding can lead to the significant compaction of DNA. However, the data are in disagreement if the compaction occurs abruptly via a first order coil-globule phase transition with a observed metastable state (Pelletier *et al.*, 2012) or gradually via a second order transition as the concentration of crowding agents increases (Cunha *et al.*, 2001).

Thus far, there are no quantitative experimental studies on how osmolality and associated changes in macromolecular crowding affect nucleoid size in living bacteria. At a qualitative level it is known that hyperosmotic shock leads to the compaction of nucleoids (Cagliero and Jin, 2013, Wu *et al.*, 2019). Here we carry out microfluidic experiments to quantitatively study the role of molecular crowders in the compaction of the *E. coli* nucleoid (i) by rapidly changing the osmolality of the growth media for steady-state growing bacteria in the mother machine device, and (ii) by squeezing individual cells in a device specially designed for such measurements. We show that *in vivo E. coli* nucleoids respond to the osmolality continuously. Close to physiological crowder concentrations the nucleoid length and width change linearly. As the crowder concentration exceeds 30% of the physiological level the compressibility significantly decreases. Also, our data show that the compressibility is strongly anisotropic being higher along the long axes of the cell and it is independent of growth conditions (slow and moderately fast growth). The latter finding indicates that the overall crowding level rather than the exact composition of crowders controls the compaction in these growth conditions. Altogether, our results lend support to the idea that differently sized molecular crowders rather than any single species are the main factor compacting the bacterial DNA within the nucleoid.

## RESULTS

### Compaction of the nucleoid under osmotic shock is anisotropic and larger than the compaction of the cytosol

Our first goal was to find a quantitative relationship between crowder concentrations/volume fractions and sizes of the nucleoids. Since altering the number of crowders in the cell is not practically possible because all macromolecules are potential crowders, one has to rely on changing the volume of the cell instead. One possibility to vary cytosolic volume, and thereby alter crowder concentration, is to change the osmolality of the growth media. By increasing the external osmolality, cytosolic water leaves the cell and crowder concentration increases, while the decrease in osmolality leads to opposite results. To vary the external osmolality and observe cellular changes in real time, we image bacteria in microfluidic mother-machine devices (Wang *et al.*, 2010, Yang *et al.*, 2018), which allowed us to quickly change media without perturbing cell imaging (Fig 1A, B). To induce osmotic shock, we changed the concentration of NaCl in chemically defined growth medium. To quantify the changes in both cytoplasmic and nucleoid sizes, the cells carried tagRFP-T and HupA-mNeonGreen labels. The former label diffusively fills the cytosol while the latter binds non-specifically to DNA (Wery *et al.*, 2001). To understand how different macromolecular crowders affect DNA compaction, we studied cells in slow and moderately fast growth conditions. The doubling times in these two conditions at 28°C are T_d_ = 226 ± 103 min and T_d_ = 95 ± 24 min, respectively. We excluded fast growth because much more complicated DNA topology and nucleoid shape in these conditions would have made the interpretation of the results ambiguous. From slow to moderately fast growth, one would expect the ribosome protein ratio in the cells to increase by 1.5-2.0 times (Bremer and Dennis, 2008, Ehrenberg *et al.*, 2013, Dai *et al.*, 2017) allowing differentiation of the effects arising from stable RNA and protein based crowders on the compaction of the nucleoid.

**Figure 1.**
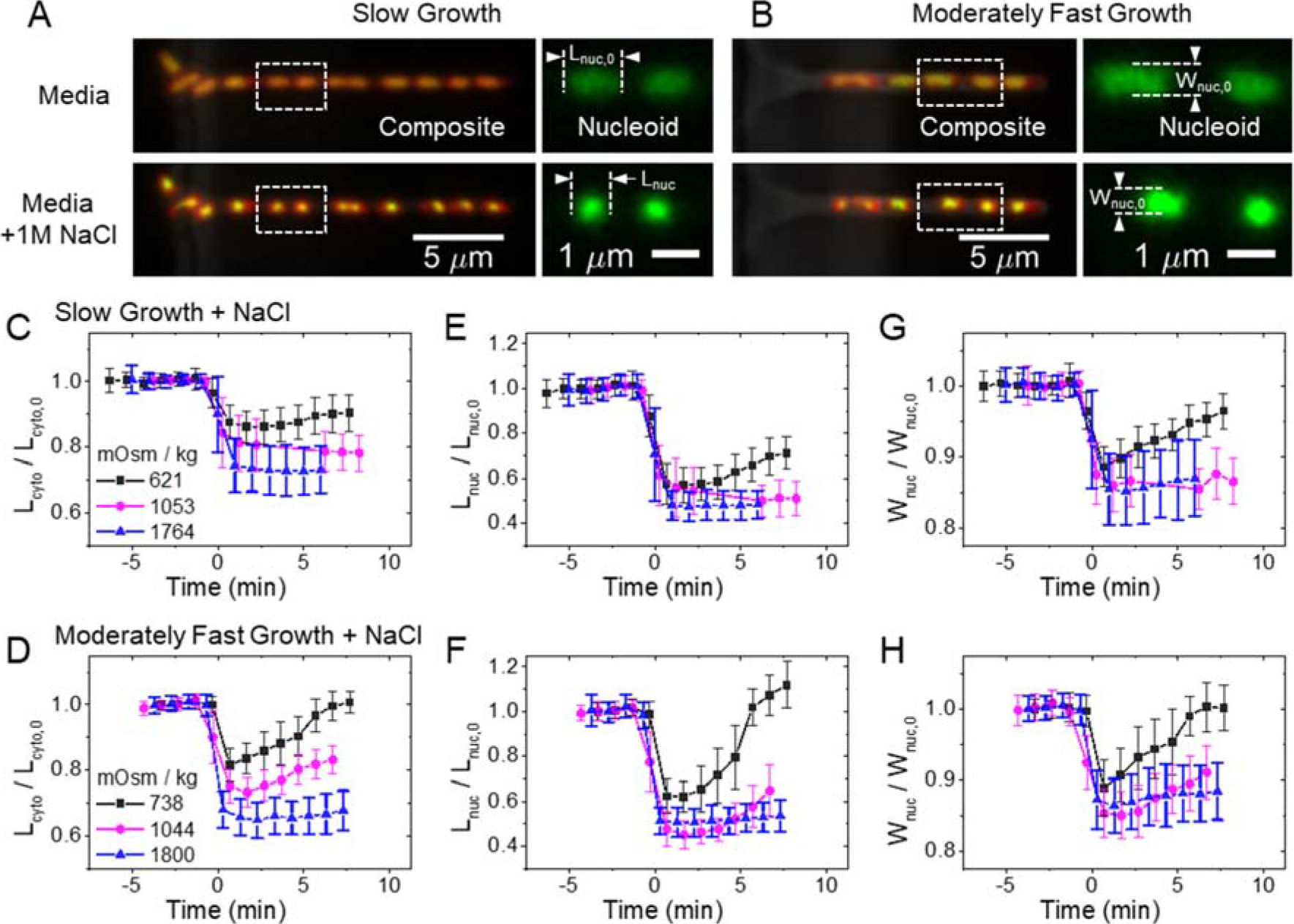
Kinetics of the cytosol and nucleoid compaction during hyperosmotic shocks. (A) Left: Images of *E. coli* cells growing in mother machine channels before (top) and 1 min after osmotic shock (bottom). The images are composites of phase contrast (grey), and fluorescence images of nucleoid (green) and cytosol (red). Right: Nucleoid images for the two cells indicated by a dashed box in the left image. The cells were grown in slow growth conditions in M9 glycerol medium. (B) The same strain in moderately fast growth conditions in M9 glucose + CAS medium. (C, D) Relative changes of cytosolic lengths for three hyperosmotic shocks in slow and moderately fast growth conditions, respectively. The shock magnitudes are indicated in lower left. *L*_cyto,0_ is cytosolic length right before the shock. The same for nucleoid length (E, F) and width (G, H). Error bars correspond to std. The number of cells analyzed in each measurement is reported in SI Table S2.

As expected, the change in external osmolality by NaCl leads to rapid changes in cell volume and nucleoid size for both growth conditions (Fig. 1A, B SI Movie M1, M2). These changes occurred in about a one-minute period (Fig. 1C-H). The rapid changes in cell shape observed here are consistent with earlier reports (Pilizota and Shaevitz, 2013, Rojas *et al.*, 2014). Interestingly, the changes in nucleoid dimensions followed the same time-dependence as changes in cell dimensions. However, dimensions of the nucleoid changed to a larger extent than the overall dimensions of the cell as will be detailed later. Although the focus of this study is not on recovery processes, in milder osmotic shocks (less than about 0.7 Osm kg^−1^) the recovery of cells is visible within 5 minutes after the shock. In these mild shocks, recovery of cell shape and nucleoid shape followed the same time-dependence consistent with the idea crowding is responsible for the nucleoid compaction.

To quantify changes in nucleoid and cell sizes at different crowding levels we analyzed cells in early stages of the cell cycle when their nucleoid has an ellipsoidal shape. In the studied growth conditions, such morphology appears in nucleoids that are less than half replicated while in later stages of replication the nucleoids obtain a characteristic bilobed shape (Bates and Kleckner, 2005, Männik *et al.*, 2016). We limit our study to single-lobed nucleoids because these can be easily characterized by their length and width measured along the long and short axes of the cell, respectively. We followed changes in these two parameters along with changes in cytoplasmic sizes in both hypo- and hyperosmotic conditions (Fig 2 A-F). However, changes in cytoplasmic volume were essentially negligible even for the most hypoosmotic shock possible when the regular growth medium was replaced by distilled water (0 Osm). Thus, almost all our studies cover hyperosmotic conditions.

**Figure 2.**
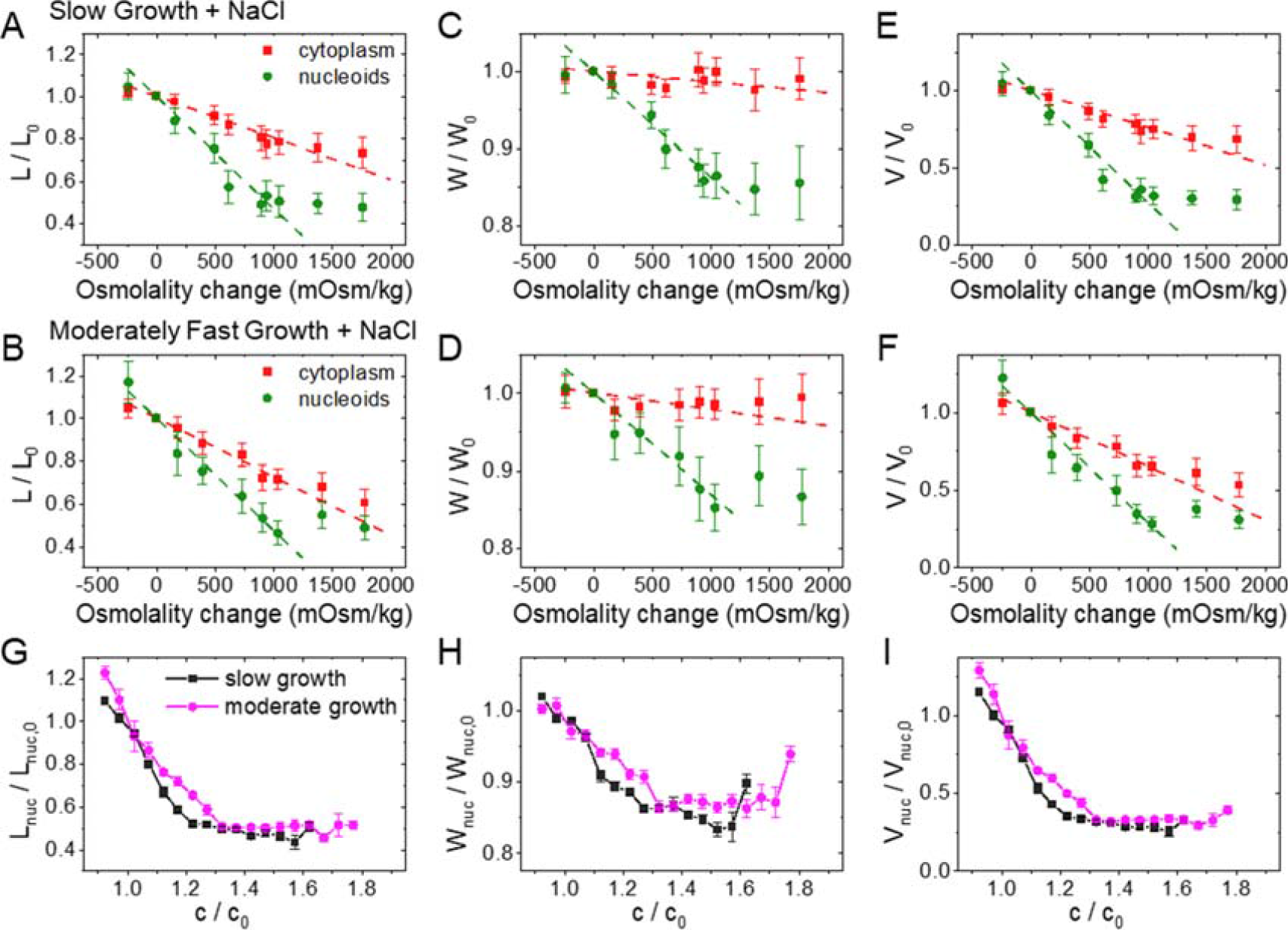
Change in nucleoid dimensions as a function of external osmolality and as a function of crowder concentration. (A, B) Relative change of nucleoid (green circles) and cytosolic (red squares) lengths at different osmotic shock in slow and moderately fast growth conditions, respectively. Each osmolality value corresponds to a separate measurement. Error bars are std. Dashed lines are linear fits to the experimental data in the range from −0.25 to 1.25 Osm kg^−1^. The best fit parameters can be found in SI Table S3. The same for nucleoid and cell widths (C, D) and calculated volumes (E, F) in these two growth conditions. (G) Nucleoid length, (H) width, and (I) calculated volume in slow (black squares) and moderately fast (magenta circles) growth conditions as a function crowder concentration. The crowder concentrations (*c*) are relative to those in normal growth medium without excess NaCl (*c*_o_). The details of calculating these concentrations can be found from SI Text: *Determination of crowder concentration based on the intensity of fluorescent reporters*. Error bars represent s.e.m.

In some cells, the hyperosmotic treatment resulted in plasmolysis. The majority of these cases showed detachment of the plasma membrane from the cell wall at the cell poles; detachments from the cylindrical parts of the cell body were less frequent, which has also been reported by others (Pilizota and Shaevitz, 2013). In cells that plasmolyzed from their side walls, the cell width is poorly defined. Moreover, in such cells nucleoids become irregularly shaped. For these reasons we excluded side-wall-plasmolyzed cells from further analyses. We found that in both moderately fast and slow growth conditions, the increase in external osmolality leads to an approximately linear decrease in cell length for osmotic shocks although above ≈1.2 Osm kg^−1^ the decrease slowed (Fig. 2A, B). Similar to cell length, the nucleoid length also decreased linearly with the osmolality change for smaller osmotic shocks (<0.9 Osm kg^−1^). At the same time the relative changes in nucleoid length were about 2.5 times as large as the relative changes in cell length (SI Table S3). At above about 0.9 Osm kg^−^1 the change of nucleoid length ceased. Nucleoid width behaved qualitatively similar to nucleoid length while the cytoplasmic width did not show significant changes throughout the range of osmolalities studied (Fig. 2 C, D). To rule out that the plateauing of nucleoid dimensions at higher osmolalities is caused by the point-spread-function of the microscope, we measured the diameters of 100 nm fluorescent beads for comparison. The measured diameters of the beads were, by more than a factor of two, smaller than the smallest measured nucleoid dimension at the highest osmotic shock (SI Fig. S1). Thus, these measurements confirmed that the measured dimensions reflect the intrinsic size of the nucleoid in the plateau region. The presence of a plateau region shows that, *in vivo*, the nucleoid undergoes a transition from a linearly high compressibility to a non-linear low compressibility regime upon compaction.

Comparison of nucleoid width and length curves showed strongly anisotropic compaction. The changes in nucleoid width at smaller osmolality changes (<0.9 Osm kg^−1^) was by a factor of 3.7 smaller than the corresponding change in the nucleoid length. The anisotropy was also present in the plateau region where the nucleoid length was compressed to 50% while its width only compressed to 70% of its original size. The nucleoid volume calculated based on these values decreased to 30% of its original size (Fig. 2 E-F). The anisotropic compaction leads to a spherical nucleoid having an aspect ratio of one at osmolalities of about 0.9 Osm kg^−1^ and above (SI Fig. S2). The spherical shape indicates underlying isotropic organization of the nucleoid.

### Nucleoid compaction is a second order phase transition and independent of growth conditions

To relate changes in osmolality to changes in crowding of the cytosolic environment we calculated crowder concentration immediately after the osmotic shock (*c*) relative to that during regular growth conditions (*c*_*0*_). This concentration change applies to any cytosolic molecular species whose diffusion across the plasma membrane during osmotic shock can be neglected. It includes not only macromolecular crowders but also all small molecules and ions. For all these molecules relative concentration change (c/c_0_) equals the inverse of their relative cytosolic volume change (V_cyto,0_/V_cyto_). However, instead of using the latter, we determined the relative concentration of cytosolic tagRFP-T label before and after salt treatment (for details see SI Text), and used this value for c/c_0_. We chose this approach because even though we only analyzed cells that did not show apparent plasmolysis from their lateral cell walls, we could have overlooked some plasmolyzed regions during visual inspection of the images. Overall, the two methods yielded comparable results at smaller osmotic shocks but deviated somewhat from each other at larger ones (SI Fig. S3).

Our data show that the length, width, and volume of the nucleoid decreased, initially, linearly as the crowder concentrations increased (Fig. 2G-I). The linear relationship corresponds to a constant compressibility of the nucleoid by the crowders. Here the compressibility, *κ*, is defined as the relative change in volume upon the change in osmotic pressure, Δ*P*, by *κ* = −(*ΔV*_*nuc*_/*V*_*nuc*_)/*ΔP*. In the lowest order approximation *P*~*c*, and in this case *κ* is proportional to −Δ*V*_*nuc*_/Δ(*c*/*c*_0_). The latter corresponds to the slope of the curves in Fig. 2I. Once the crowding level exceeded about 30% of the level in normal growth conditions, the compressibility decreased sharply. The transition to the low compressibility regime was smooth unlike in several previous *in vitro* studies (Pelletier *et al.*, 2012, Krotova *et al.*, 2010, Yoshikawa *et al.*, 2010). There was no sign of co-existence of collapsed and extended DNA conformations during visual inspection of the nucleoid images nor in the raw data of nucleoid lengths and widths (SI Fig. S4). Both findings together indicate that coil-globule transition of the nucleoid is a second rather than first order phase transition *in vivo* conditions.

Interestingly, we found that the dependence of the nucleoid size on the crowder concentration is almost the same in both studied growth conditions (Fig. 2G-I). This outcome is surprising because concentrations of the main macromolecular crowders, the ribosomes, and the proteins, are expected to be significantly different in these two growth conditions (Ehrenberg *et al.*, 2013, Dai *et al.*, 2017).

### Mechanical squeezing measurements confirm osmotic shock data

In addition to removing cellular water content via osmosis we also carried out measurements where we removed water by mechanical squeezing of cells. For that purpose we constructed a microfluidic device, which we refer to as the microanvil (Fig. 3A, for details see SI Text). This device allowed us to mechanically press out part of the cytosolic content while imaging the cell (Fig. 3B). Our first proof-of-principle devices had the smallest anvil dimension of about 2 μm. In future designs this dimension can be reduced. Since 2 μm constitutes a sizeable portion of cell length (2-4 μm), we studied longer than normal cells. The cells were elongated by inducing SulA expression from an extra plasmid copy, thereby inhibiting cell division (Dajkovic *et al.*, 2008). The elongated cells had lengths in the range of 7-15 μm and had multiple nucleoids. The amount of cytosolic content that can be removed from the cells can be controlled by an external pressure applied by the anvil. However, at higher pressures the anvil divides the cell into two distinct compartments (Fig. 3B, SI Movie M3). In the region between the two compartments the cytosolic content of the cell is almost completely removed (SI Fig. S5). Further increases to the externally applied pressure by the anvil do not appreciably change the cytosolic volume of the cell. For this reason, the maximum volume that was removed was limited to about 25% of the initial cytosolic volume in these experiments.

**Figure 3.**
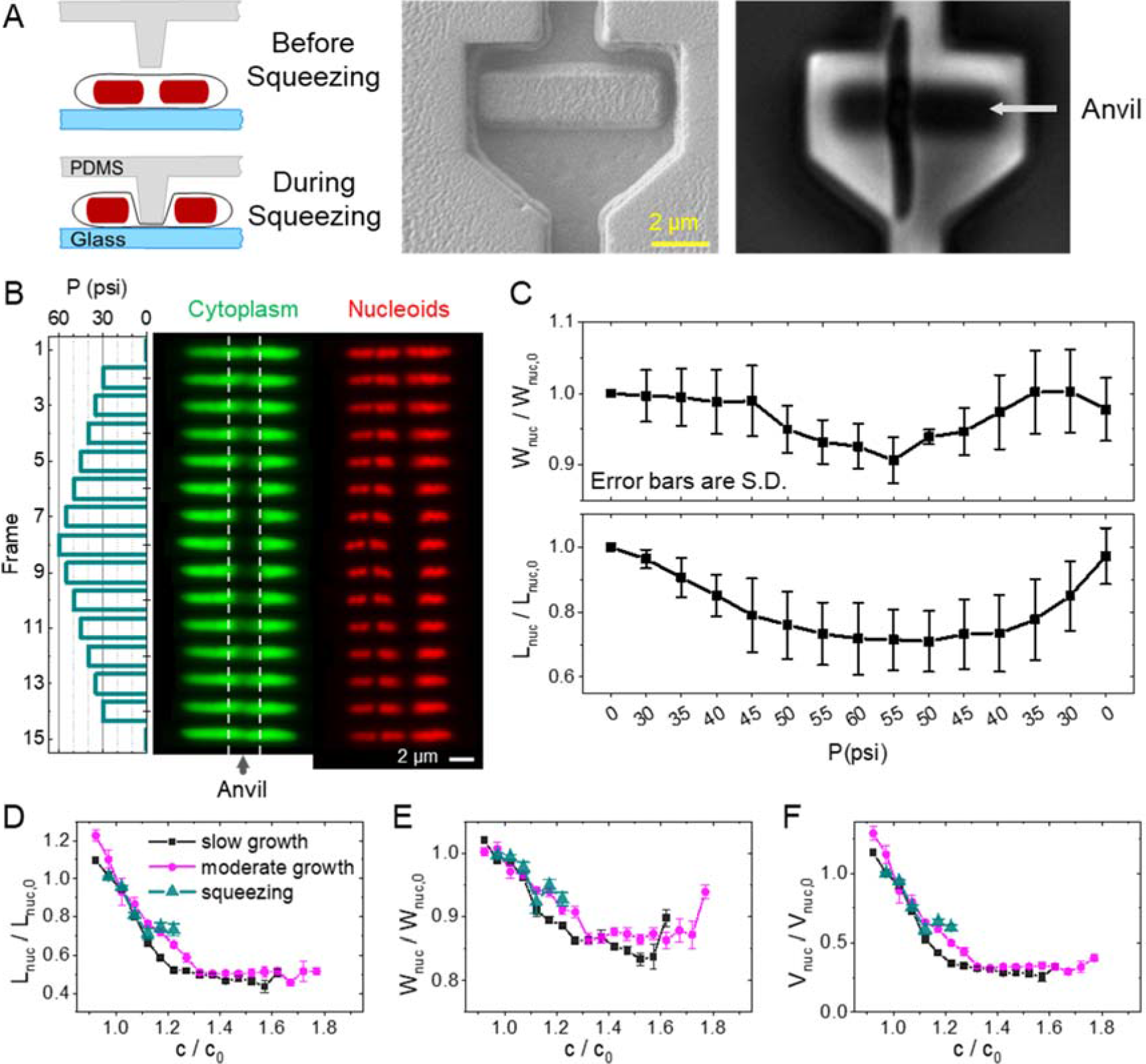
Measurements of nucleoid compaction during cell squeezing. (A) Left: Schematics of the measurement. A pressure actuated valve is fabricated with a small anvil-like protrusion. Externally applied pressure deflects the anvil downward and squeezes about a 2 μm long portion of the underlying cell. Middle: SEM image of valve and anvil. Right: phase image of cell trapped under the anvil. (B) Response of an elongated *E. coli* cell to a pressure cycle. Left panel: Change of externally applied pressure during the measurements. In the 1^st^ (bottom) and the last (top) frame the cell is not squeezed. Middle panel: images of cytoplasmic mNG label during this pressure cycle. The region between dashed lines correspond to the portion of the cell where the anvil touches it. Right panel: Nucleoid images for this cell. (C) The average nucleoid widths (top) and length (bottom) for this cell. The average is over three nucleoids in this cell and error bars reflect std. Both the width and length of the nucleoids are relative to those at the beginning of the squeezing cycle (W_nuc,o_ , L_nuc,o_, respectively). (D) Comparison of relative nucleoid length as a function of crowder concentration from squeezing measurements (green triangles) to that from osmotic shock measurements (magenta circles and black squares). The same for nucleoid width (E) and calculated volume (F).

Compaction of the nucleoid increased during stepwise pressure ramps and recovered when the pressure was lowered (Fig. 3C). As with measurements with osmolality variation, efflux and influx of water and other cytosolic components were taking place at a faster time scale than the measurement rate (0.5 min^−1^). After one pressure cycle, the majority of cells were capable of resuming growth.

We determined nucleoid dimensions and volumes from these measurements as a function of the changes in crowder concentrations and compared these results with the findings from the osmotic shock measurements (Fig. 3D-F, S6). Within uncertainties of the measurement, the two approaches yielded indistinguishable results thus further confirming the osmotic shock measurements.

### Transertion linkages and polysomes do not control nucleoid size in slow growth conditions

We next aimed to understand which of the macromolecular crowders in the bacterial cytosol play the dominant role in compaction of the nucleoid. Several lines of research point out that the main compacting agents among different macromolecular crowders species are poly-ribosomes (polysomes) (Joyeux, 2016, Mondal *et al.*, 2011, Bakshi *et al.*, 2014, Bakshi *et al.*, 2015). This conclusion has been partially drawn from studies of cells treated with rifampicin, a transcription halting drug. The drug appears to affect the nucleoid size via two different mechanisms in fast growth conditions. At shorter time scales (0-5 min) nucleoids compact (Bakshi *et al.*, 2014). The compaction has been explained to be the result of severing transertional linkages between DNA and the inner membrane. These linkages are expected to keep the nucleoid in an expanded state and to resist the compaction effects from the crowders. In longer timescales (>10-15 min), rifampicin leads to the expansion of chromosomal DNA so that it appears to fill the whole cytosolic volume (Bakshi *et al.*, 2015). The expansion has been interpreted as the result of polysomes dissociating into 30S and 50S ribosomal subunits and by the lower ability of these subunits to compact the nucleoid (Bakshi *et al.*, 2015). Since the dissociation of the polysomes during rifampicin treatment has such drastic effects on the nucleoid size, one can expect them to be the dominant species in compacting the nucleoid.

We were interested in understanding if the above described nucleoid behaviors during rifampicin treatment also hold in moderately fast and slow growth conditions where ribosome and polysome concentrations are expected to be lower (Ehrenberg *et al.*, 2013, Dai *et al.*, 2017). Treating moderately fast growing cells with rifampicin, we observed an initial compaction of the nucleoid at about 5 min timescale (Fig 4 A-C) consistent with the earlier report (Bakshi *et al.*, 2014). Although the effect of compaction in our experiments was smaller (3-5%) (Fig. 4 A-C insets) than in the previous report (about 8%), which was carried out in fast growth conditions, the presence of nucleoid contraction supports the idea of the existence of transertion linkages. However, their effect in determining the size of the nucleoid appears rather modest in this growth condition. At timescales longer than 5 min we observed the expansion of the nucleoid (Fig. 4 A-C). This finding is also consistent with the earlier hypothesis that ribosome subunits are less effective in compacting the nucleoid than the assembled polysomes (Bakshi *et al.*, 2014, Bakshi *et al.*, 2015, Bakshi *et al.*, 2012).

**Figure 4.**
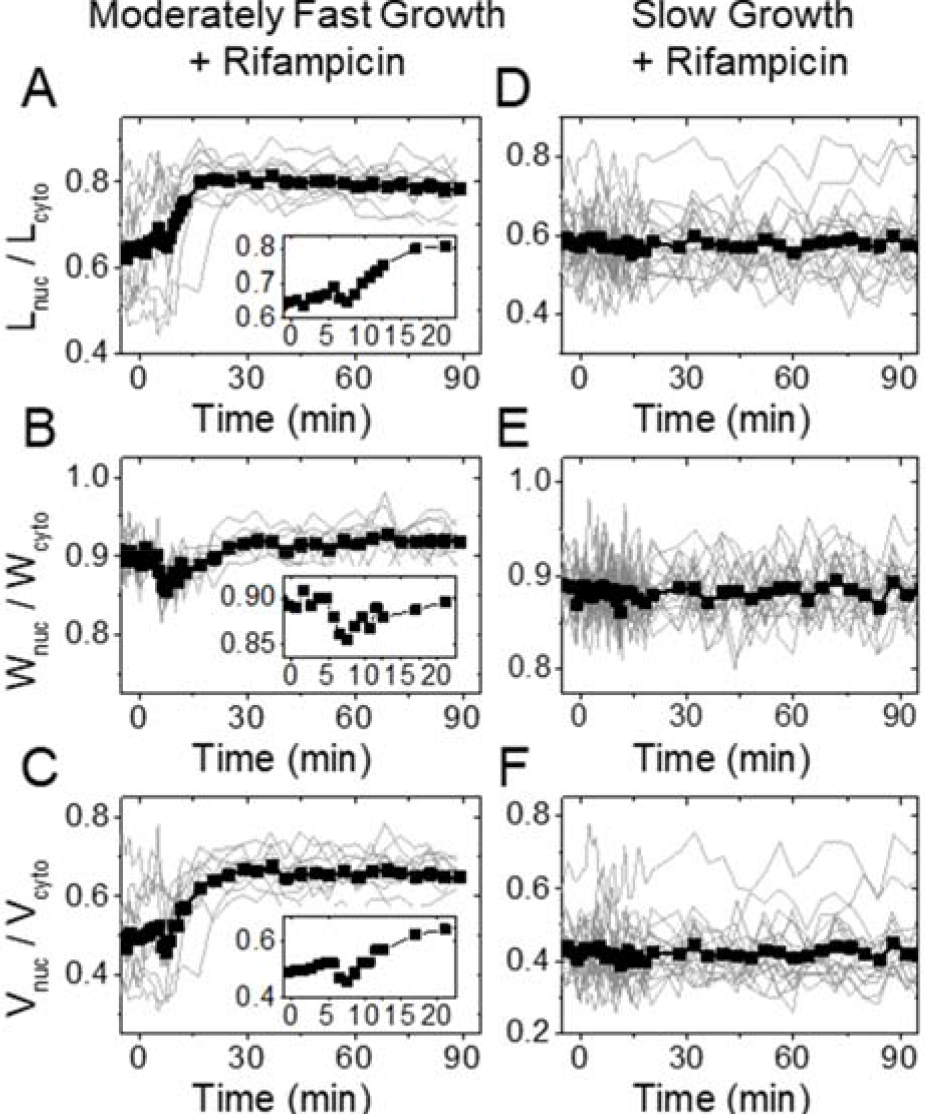
Change in nucleoid dimensions under rifampicin treatment in moderately fast (left) and slow (right) growth conditions. 300 μg ml^−1^ rifampicin is administered to cells at 0 minutes. There is 2-3 min delay in fluidic lines before the drug reaches the cells. (A, D) Ratio of nucleoid to cell length as a function of time. Traces from individual cells are shown by thin lines, and the population average trace by a thick line with squares. N=20 for slow and N=11 for moderately fast growth conditions. Inset in moderately fast growth conditions is zoomed in from the beginning of the trace. (B, E) The same for nucleoid width. (C, F) The same for the calculated nucleoid volume.

Interestingly, all the above changes in nucleoid dimensions were absent in slow growth conditions (Fig. 4 D-F) although rifampicin manifested itself in stopping cell growth (SI Fig. S7). Lack of change in nucleoid dimensions at short time scales (< 5 min) show that the effect of transertion in determining nucleoid size is completely negligible in slow growth conditions. This is perhaps expected given an overall slower protein synthesis rate and therefore a smaller number of transertion linkages in this growth condition. By the same token, a smaller concentration of ribosomes and polysomes in slow growth conditions can help explain the lack of nucleoid expansion at longer time scales (>5 min). However, the latter leads to a further inference that polysomes and ribosomes have a small contribution to the compaction of the nucleoid in slow growth conditions.

### Limited role of polysomes in compacting the nucleoid in moderately fast growth conditions

To further understand the role of polysomes in the compaction of the nucleoid, we applied osmotic shocks to cells that were treated for 20-25 min with rifampicin (Fig. 5). During this treatment period, the majority of the polysomes should dissociate into 30S and 50S subunits given that mRNA lifetime in *E. coli* is about 5 min (Bernstein *et al.*, 2002). The latter assessment is also consistent with the observed nucleoid expansion at the 15 min time scale in moderately fast growth rates in our measurements (cf. Fig. 4 A-C). Note that the chosen 20-25 min period is short enough for the concentrations of protein and stable RNA species in the cytosol to not alter significantly. We estimate their change to be less than 10% due to their dilution by residual cell growth while their decrease due to degradation can be expected to be minimal during this period. If ribosomal subunits are less effective in compacting the chromosomes (Bakshi *et al.*, 2014, Bakshi *et al.*, 2015, Bakshi *et al.*, 2012) then we would expect weaker compaction of nucleoids in rifampicin treated cells in osmotic shock measurements. Contrary to this expectation, the nucleoid compaction was somewhat larger in rifampicin treated cells in both growth conditions (Fig. 5). In moderately fast growth conditions, a slight increase in compaction could be assigned to the fact that prior to the osmotic shock, the nucleoid was larger in rifampicin-treated cells than in untreated ones. To compensate this effect, we extrapolated nucleoid lengths from the beginning of the treatment to the point where salt shock occurred, assuming an increase in nucleoid length would have followed the same increase as during normal growth (SI Fig. S8). The resulting curves still showed negligible effects from rifampicin treatment on the nucleoid compaction curves (SI Fig. S9). Furthermore, we compared nucleoid width distributions in addition to their normalized distributions. We found the former to be indistinguishable for rifampicin treated and untreated cells (SI Fig. S10) showing that the nucleoid compacted into the same final width as a result of hyperosmotic shock irrespective of prior rifampicin treatment. Thus, these results show that rifampicin does not alter nucleoid compaction during hyperosmotic shock even if it leads to nucleoid expansion at higher growth rates. If the assumption that rifampicin dissociates polysomes is correct, then these data show that polysomes do not significantly contribute to nucleoid compaction during osmotic shocks.

**Figure 5.**
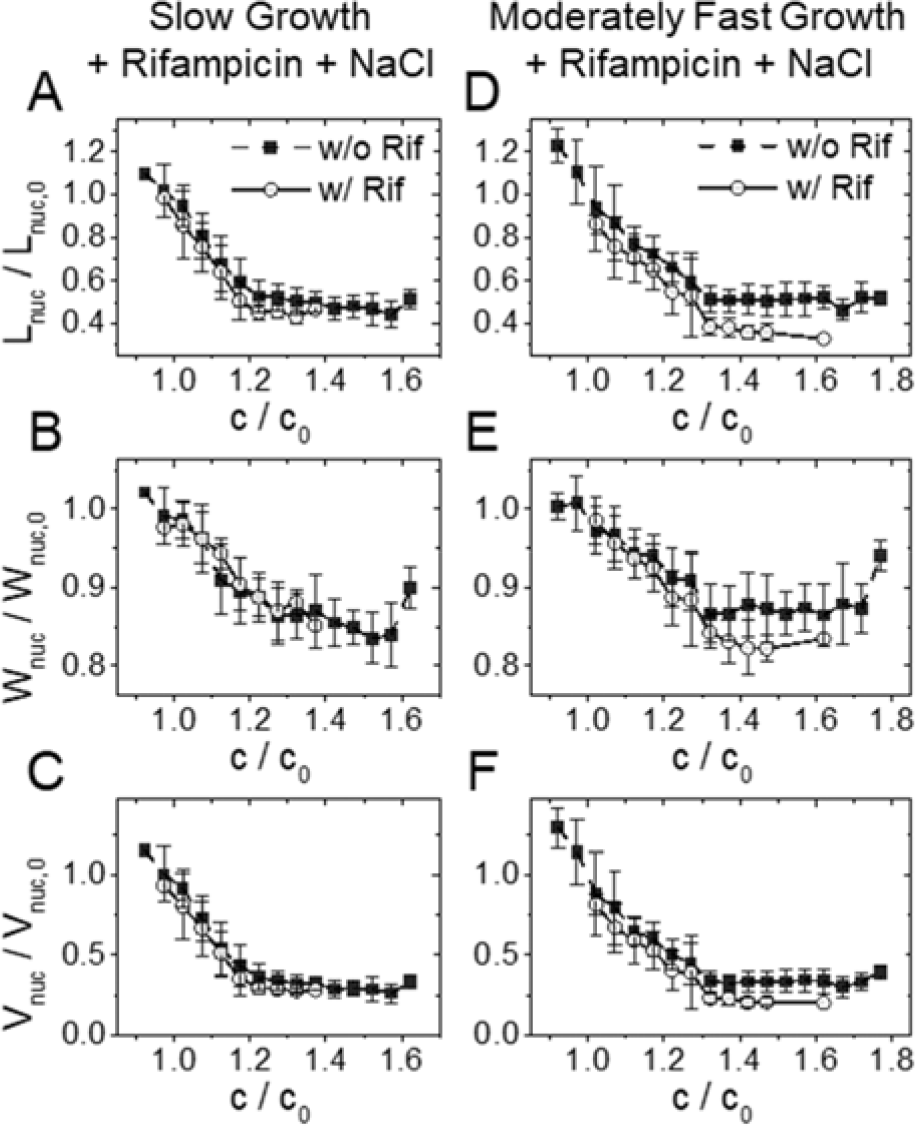
Effect of osmotic shock after rifampicin treatment. Cells were first treated for 25-30 min with 300 μg ml^−1^ rifampicin and then the concentration of NaCl in the medium was changed. Rifampicin was also present in the medium during osmotic shock. Nucleoid dimension right after osmotic shock divided by the same dimension right before the shock (solid line open circle). For comparison the same ratio without rifampicin treatment is also shown (black dashed line solid square; from Fig. 2G-F). (A, B) Relative nucleoid length as a function of relative crowder concentration. (C, D) The same for relative nucleoid width, and (E, F) for the calculated volume.

### Crowding model qualitatively agrees with experimental findings

We sought to explain the measured data using Brownian dynamic simulations of chromosomes in crowded and confined environments. Our computational approach is similar to the one reported by Kim et al (Kim *et al.*, 2015) (for more details see SI Text). Extending the previous report, we also calculate nucleoid width and volume in addition to nucleoid length to compare these quantities to experimental values. The model represents the chromosome as a chain of linked beads and crowders as smaller-sized unlinked beads. Both species interact via repulsive excluded volume interactions and are confined to a cylindrical volume (Fig. 6A). The model chromosome corresponds to a single fully replicated circular *E. coli* chromosome.

**Figure 6.**
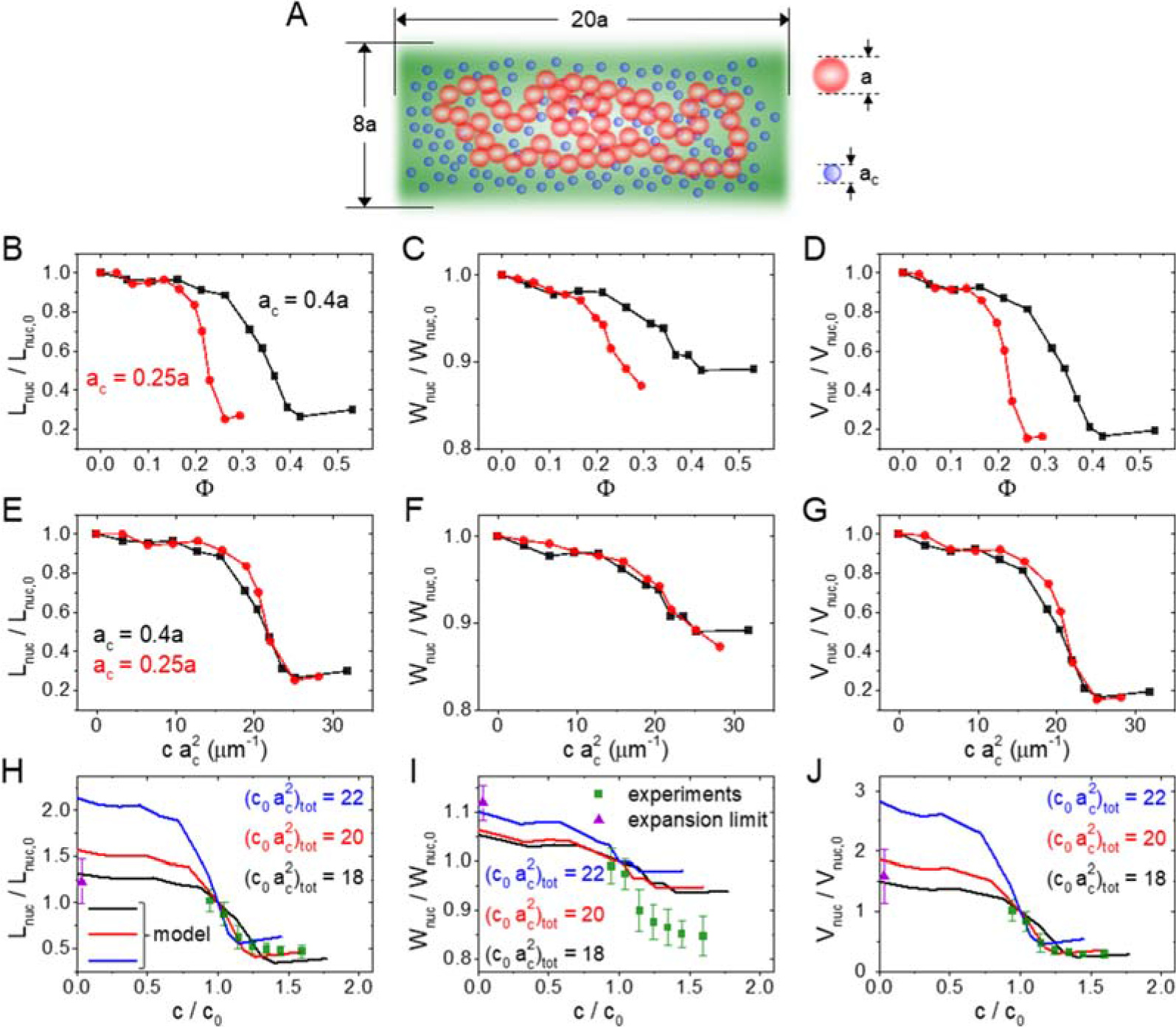
Comparing a polymer model to the experimental data. (A) A model of DNA consists of 150 beads with diameter *a* that are confined to a cylindrical volume. The same volume is also occupied by a variable number of crowders having diameters *ac*. For details of the model see SI Text: *Coarse-grained Brownian Dynamics simulations*. (B-D) Calculated nucleoid length, width, and volume for two different size crowders as a function of their volume fraction. (E-G) Calculated nucleoid length, width, and volume as a function of effective “crowding level” defined as 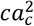. The curves with different crowder diameters collapse into a single one. (H-J) Comparing modelled nucleoid length, width, and volume (solid lines) to experimental data from slow growth conditions (green squares; from Fig. 2G-F). The experimental data is augmented by a data point at the zero crowder concentration limit where the nucleoid is expected to extend over the whole cytosolic volume (violet triangle). A set of curves from the model with “crowding level” 18, 20, 22 μm^−1^ are plotted. Of those the best agreement between the experiment and the model is at 20 μm^−1^. The error bars for the data are std.

According to model calculations the length, width, and volume show approximately sigmoidal dependence on the volume fraction of crowders (Φ) or the concentration of crowders (Fig. 6B-D, SI Fig. S11). The former is defined as an effective total volume of crowders divided by the volume of the cytoplasm and it is proportional to the concentration of crowders. Similar dependence for nucleoid length on the volume fraction was also observed in earlier models (Shendruk *et al.*, 2015, Kim *et al.*, 2015). The model predicts that in both low and high crowder volume fractions nucleoid dimensions are insensitive to the amount of crowders in cellular volume. The transition between the two regimes occurs continuously, i.e. the coil-globule transition in this model is second order. All these predictions are consistent with our experimental data on a qualitative level.

We calculate the aforementioned dependences for two different crowder sizes. The model predicts that the mid-point of nucleoid compaction occurs at larger volume fractions for larger crowders. However, a smaller concentration of larger crowders is needed to compact the nucleoid to the same level (SI Fig. S11). Kim *et al.* have previously found that multiplying crowder concentration by the square of the crowder diameter, i.e. by 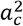, collapses 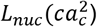 for different size crowders to a single curve (Kim *et al.*, 2015). Our model indicated that this scaling applied in addition to the nucleoid length also to its width and volume (Fig. 6 E-G). Furthermore, results by Kim *et al.* showed that for a polydisperse ensemble of crowders *L_nuc_* vs 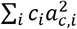 curves collapsed for different crowder mixtures into a single curve (Kim *et al.*, 2015). The above sum is taken over all crowder species. Thus, the level of crowding in polydisperse samples is, according to the model, characterized by a single parameter 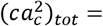 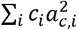 which one could calculate knowing the concentrations and diameters of all crowders.

We used the scaling behavior of model curves to compare them to our experimental data. By adjusting the crowding parameter 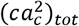, which corresponds to the crowding level in the regular medium, we found the best match between experiment and model for 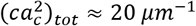 (Fig. 6 H-J). In this comparison we also accounted that at zero crowder concentration the nucleoid must fill the whole cell volume. This extra constraint was determined as the ratio of the measured cytoplasmic size to nucleoid size in regular growth medium (violet triangles in Fig. 6H-J). As can be seen from Fig. 6H, the model predicts the experimentally observed length dependence on the crowding level reasonably well. The agreement is poorer for the nucleoid width, where the experiment shows much larger variation and thus smaller anisotropy in the compressibility (Fig. 6I). Taking the coarse-grained nature of the model, some quantitative discrepancies can be expected; yet the main experimental characteristics are clearly represented in the model.

## DISCUSSION

### Smooth coil-globule transition of nucleoids in the cellular environment

Our measurements show that under hyperosmotic treatment and during mechanical squeezing *E. coli* nucleoids undergo rapid compaction on a timescale less than one minute. The compaction occurs concurrent with the changes in cytoplasmic volume, but during mild shocks the nucleoid volume decreases by a factor of about 2.5 times more than the cytoplasm. The shrinkage of the nucleoid can be explained by an increased osmotic pressure that acts on it when water leaves the cell. As our squeezing measurements show, the increased osmotic pressure arises from elevated concentrations of macromolecular crowders rather than from increased concentrations of some small molecules and ions. These conclusions are in overall agreement with previous *in vitro* studies where artificial crowding agents or ions have been used. However, several *in vitro* studies have reported an abrupt transition from an extended state of the liberated nucleoid (coil) to a highly compacted state (globule) (Krotova *et al.*, 2010, Pelletier *et al.*, 2012, Yoshikawa *et al.*, 2010). Here we find that *in vivo* the compaction of the nucleoid is a continuous function of cytoplasmic crowder concentration. Therefore, the associated coil-globule transition of the chromosome is the second rather than the first order phase transition in the cellular environment. The difference between *in vivo* and *in vitro* experiments can be expected taken the different nature of crowders and different organization of DNA due to DNA binding proteins and supercoiling. The physiological implication of the second order transition is that the nucleoid and cellular processes related to the nucleoid respond to osmolality changes continuously, instead of maintaining a steady nucleoid homeostasis up to a certain shock magnitude and then completely losing this homeostasis in the fashion of an on-off switch.

### Anisotropic compressibility of the nucleoid and bottlebrush-like organization of the chromosome

Our data show that the nucleoid compresses anisotropically at lower crowder concentrations. Its longitudinal compressibility is about four times as high as their radial one. At osmotic shocks of about 1 Osm kg^−1^ nucleoids become spherical and remain spherical at higher shocks (SI Fig. S2 C, D). Based on EM images of lysed cells (Kavenoff and Bowen, 1976), it has been proposed that *E. coli* nucleoids have a bottlebrush-like organization with supercoiled segments or just DNA loops stretching out radially from a backbone, which is aligned with the long axes of the cell (Wang *et al.*, 2013). This view also has some support from 3C/Hi-C studies of *Caulobacter crescentus* (Le *et al.*, 2013), *Bacillus subtilis* (Marbouty *et al.*, 2015), and *E. coli* (Lioy *et al.*, 2018) chromosomes. These studies all show well-defined chromosomal interaction domains, which could correspond to supercoiled segments that stretch radially out from a common backbone.

It can be expected that bottlebrush-like organization leads to anisotropic compressibility. It should be harder to compact plectonemic supercoils along their length, which according to this model are oriented radially relative to the long axes of the cell. At the same time spacing between supercoils allows them to be easily compacted along the cell length. Although our data appears consistent with the above explanation, the modeling results show that anisotropic compaction can be expected even for the chromosome that lacks supercoiling and is a consequence of cylindrical confinement of the chromosome by the inner membrane. Moreover, it would be unlikely that the anisotropic bottlebrush-like chromosome would be compacted to a spherical entity. Bottlebrush like structure should retain its anisotropy, and as a result, its aspect ratio should differ from one, which is contrary to what is found in our experiments (SI Fig S2 C, D). Altogether, our data favors a more disordered organization of plectonemic supercoils than envisioned by the bottlebrush model where supercoils emanate from a single linear backbone.

### Interplay of different crowders in compacting the nucleoid

Our data provides new information on how different cytoplasmic macromolecular crowders affect nucleoid compaction. In particular, we find that polysomes cannot be the sole dominant crowder species that leads to nucleoid compaction despite their large volume fraction, high charge state and prominent exclusion from the nucleoid. These conclusions are based on essentially identical nucleoid compaction curves in two different growth rates, and insensitivity of these curves to rifampicin treatment. In the following discussions, in order to further rationalize these findings, we estimate the contribution of different macromolecular crowders in compacting the nucleoid based on their literature reported abundances and sizes. The crowder groups that we consider are cytoplasmic proteins, 30S and 50S ribosomal subunits, tRNA, and poly-ribosomes (polysomes). Not all cytoplasmic proteins qualify as crowders. We exclude DNA binding proteins from this group. Also, we group proteins that are involved in translation together with their rRNA and tRNA counterparts. There are an estimated 3 million proteins in *E. coli* in fast growth conditions (Milo, 2013). Of those, 20-25% qualify as cytoplasmic crowders that are not part of ribosomes, chromosomes, and envelope layers. In slow growth, where the cell needs to synthesize metabolic components from simpler molecules, the fraction of cytosolic crowders can increase to 40% of the total proteome (Li *et al.*, 2014). 80-85% of the cellular RNA content should be a part of the actively translating ribosomes. These ribosomes form polysomes. The remaining 15% of the RNA mass should be in the form of 30S and 50S subunits in moderately fast growth conditions (Dai *et al.*, 2017). In slow growth conditions the corresponding numbers are 65% for polysomes and 35% for subunits. tRNA abundance is about nine molecules per one ribosome (Bremer and Dennis, 2008). In slow growth conditions, ribosome to protein mass ratio is estimated to be 1.5-2.0 times smaller than that in moderately fast growth conditions (Ehrenberg *et al.*, 2013). This appears to be the result of a decrease in ribosome concentration in slow growth while its protein concentration remains approximately unchanged.

The results from Kim *et al.* (Kim *et al.*, 2015) and from our modelling predict that different crowders contribute to the nucleoid compaction not by their volume fraction but via their crowding level given by 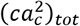. Using the above numbers (SI Table S4) we estimate that in moderately fast growth conditions polysome and protein based crowders are the main contributors (44%, and 35%, respectively) while the contributions from rRNA subunits and tRNA are smaller (15% and 6%, respectively) (SI Table S5). The large contribution of polysomes stems from their large (excluded) volume, which also leads them to be spatially separated from the nucleoid; even at small concentrations. Large contributions of proteins arise from their much larger numbers. Unlike ribosomes, the fraction of proteins that are excluded from the nucleoid needs to not be very large. Even a small fractional difference (1-5%) in protein concentration between the nucleoid and cytosolic phase is sufficient to cause a similar effect on nucleoid compaction than the ribosomes because of their much larger numbers and translational entropy. Altogether, polysomes and protein crowders are distributed very differently between cytosolic and nucleoid phases but they contribute comparably to nucleoid compaction in this growth condition.

Using 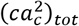 again as a metric, we estimate that in slow growth conditions proteins contribute 67% and polysomes 20% to the total crowding level (SI Table S5). The overall crowding levels in these two growth conditions are comparable (4% higher in slow growth). Based on these arguments, comparable compaction of the nucleoids in both growth conditions is expected during hyperosmotic shock as is observed in the experiments. These estimates predict that as the growth rate slows, the dominant contribution from polysomes on nucleoid compaction changes to a dominant contribution by the proteins instead.

Clearly, the above estimates contain significant uncertainties, as they are based on limited and scattered quantitative data from the concentrations of cellular crowders and their dimensions. Despite these uncertainties, the above analysis points out that several main groups of cytoplasmic crowders, beyond polysomes, can be expected to have a significant contribution to nucleoid compaction. This broader conclusion is consistent with our experimental data.

### Physiological consequences of nucleoid size

Our data show that macromolecular crowders maintain the nucleoid in a distinct state/phase in *E. coli* where the spatial extent is sensitive to the variation in crowding levels. This state is limited to a relatively narrow level of crowding (Fig. 7). As predicted by our modeling, a 25% decrease of crowding level can lead to a completely diffuse chromosome, which fills the whole cytoplasmic volume. This scenario appears in some bacteria. It has been reported, that in *C. crescentus* the chromosomal DNA extends throughout the cytosolic volume (Llopis *et al.*, 2010); although it may be excluded from the immediate vicinity of inner membrane (Woldringh, 2010). At the same time, a 30% increase in crowding levels leads to a highly compressed nucleoid according to both experimental and modeling results. Such a nucleoid state would possibly hinder some DNA transactions, such as replication and transcription, as the unwinding of DNA strands requires more energy in the compacted nucleoid. It remains an intriguing question for further research to understand if it is a co-incidence that crowding levels are just right to maintain the nucleoid in a sensitive part of the compaction curve or is there some dedicated regulatory mechanisms involved.

**Figure 7.**
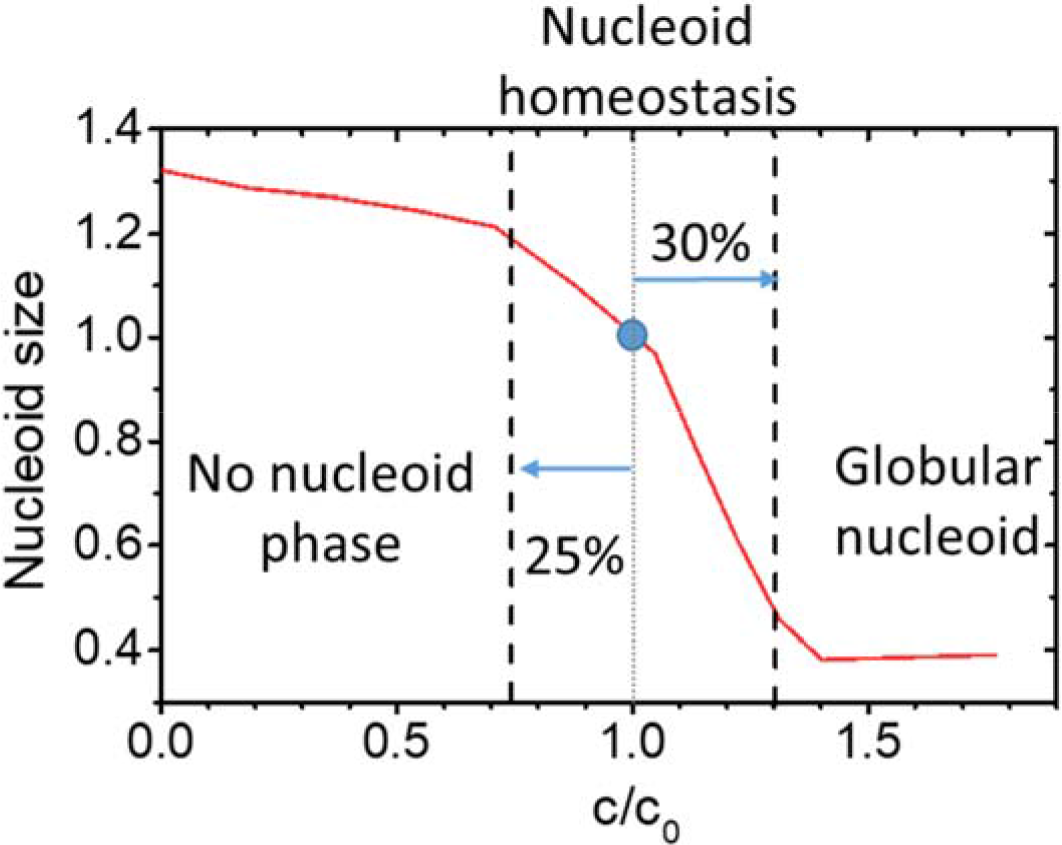
Nucleoid homeostasis in the presence of crowders. Distinct nucleoid where DNA is moderately compacted exists in a limited range of crowding levels. 25% reduction in crowding levels leads to abolishment of distinct nucleoid phase from which crowders to some degree are excluded. At the same time, a 35% increase in crowder concentration leads to the formation of globular nucleoid with high compressibility and likely of limited functionality. Red curve is model data at 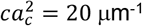 matched approximately to experimentally measured nucleoid length. The blue circle corresponds to a nucleoid size in physiological conditions.

## EXPERIMENTAL PROCEDURES

### Strains and growth conditions

All strains in this study were derivatives of *E. coli* K-12 MG1655 (SI Table S1). The genomic insertions were constructed using λ-Red recombination and shuffled between strains using P1 transduction as described in (Datsenko and Wanner, 2000). The details of strain construction can be found in SI Text. The bacteria were grown and imaged at 28°C. In slow-growth conditions cells were cultivated in M9 minimal medium (Teknova Inc., CA) supplemented with 2 mM magnesium sulfate (MgSO_4_) (Millipore Sigma, MO), 0.3% glycerol (Fisher Scientific), and 100 μg ml^−1^ leucine (Fisher Scientific). In moderately fast growth conditions M9 minimal medium was supplemented with 2 mM MgSO_4_, 0.5% glucose (Millipore Sigma, MO), and 0.2% casamino acids (Fisher Scientific). In all experiments, except squeezing measurements, cells were first plated and grown on M9 agar plates supplemented with 2 mM MgSO_4_ and 0.5% glucose. From the plate a single colony was inoculated into 2.5 ml of media. The culture was grown overnight and then injected into the microfluidic chips.

In squeezing experiments (strain DY3), cells were first plated and grown on LB agar plates supplemented with 20 μg ml^−1^ chloramphenicol and 0.2% glucose. A single colony was inoculated into M9 supplemented with 2 mM MgSO_4_, 0.5% glucose, and 20 μg ml^−1^ chloramphenicol, then grown overnight. The overnight culture was diluted to an OD600 ~ 0.002 in fresh M9 medium supplemented with 2 mM MgSO_4_, 0.3% glycerol, and 50 nM IPTG to induce expression of the extra-chromosomal *sulA* gene under the control of the *lac*-promoter. After six hours, the cells were filamentous and contained multiple nucleoids.

### Microscopy

A Nikon Ti-E inverted epifluorescence microscope (Nikon Instruments, Japan) with a 100X (NA = 1.45) oil immersion phase contrast objective (Nikon Instruments, Japan), was used for imaging the bacteria. Images were captured on an iXon DU897 EMCCD camera (Andor Technology, Ireland) and recorded using NIS-Elements software (Nikon Instruments, Japan). Fluorophores were excited by a 200W Hg lamp through an ND8 neutral density filter. A Chroma 41004 filtercube was used for capturing mCherry and tag-RFP-T images, and a Chroma 41001 (Chroma Technology Corp., VT) for mNeonGreen images. A motorized stage and a perfect focus system were utilized throughout time-lapse imaging.

### Image analysis

Image analysis was carried out using Matlab (MathWorks, MA) scripts based on Matlab Image Analysis Toolbox, Optimization Toolbox, and DipImage Toolbox (https://www.diplib.org/). For each cell analyzed, a segment connecting its two poles in the image of the cytoplasmic label was manually identified. Then, this segment defined the longitudinal axis of the cell. The segment was then broadened to nine pixels along the short axes of the cell and it was used to extract intensity line profiles for the cytoplasmic and for the nucleoid labels. Subsequently, these profiles were smoothed by least square fitting them to the exponential power function. The latter is a type of generalized Gaussian function. From these smoothened curves, inflection points were determined. The distance between the inflection points of respective profiles were used as an estimate for the cell and nucleoid lengths.

To determine the cytoplasmic and nucleoid widths intensity line profiles were generated perpendicular to the long axes of the cell. These intensity profiles, taken one pixel apart, were each fitted to a (regular) Gaussian function. From the fittings, the mean distance between the inflection points for each profile was determined. Based on earlier modeling results (Männik *et al.*, 2009), the widths were estimated as the distances between the inflection points of the Gaussians to which an additional constant offset of 80 nm was added. Finally, a mean width was calculated from averaging widths over the length of the cell.

In the microanvil measurements, cells had multiple nucleoids. To determine nucleoid lengths, the nucleoids images were first resampled using a nine-pixel-wide polyline. An averaged intensity line profile based on this polyline was then calculated. This intensity profile was then fit to multipeak exponential power functions. Each nucleoid lobe was represented by a single exponential power function. From the exponential power functions, distances between the second inflection points were determined. The latter served as an estimate for the nucleoid length. Cell and nucleoid volumes were calculated assuming they were cylinders capped with half spheres on each pole. Cell and nucleoid aspect ratios were length to width ratios.

Details of microchip design, fabrication, and usage, and coarse-grained modeling are given in the SI Text.

## ACKNOWLEDGEMENT

The authors thank Fabai Wu, Cees Dekker, and Rodrigo Reyes-Lamothe for strains and plasmids, and Sriram Tiruvadi Krishnan, Bryant E. Walker, James Weisshaar, Conrad Woldringh, and Arieh Zaritsky for valuable comments. Authors acknowledge technical assistance and material support from the Center for Environmental Biotechnology at the University of Tennessee. This work was supported by National Science Foundation research grant [MCB-1252890], US-Israel Binational Science Foundation research grant [2017004], and National Institutes of Health award under [R01GM127413]. Part of this research was conducted at the Center for Nanophase Materials Sciences, which is sponsored at Oak Ridge National Laboratory by the Scientific User Facilities Division, Office of Basic Energy Sciences, U.S. Department of Energy. A part of this research is based upon work performed using computational resources supported by the University of Tennessee and Oak Ridge National Laboratory Joint Institute for Computational Sciences.

