## Supporting Information for "The effects of polydisperse crowders on the compaction of the *Escherichia coli* nucleoid"

**This PDF file includes:**

SI Text
Figures S1 to S11
Tables S1 to S5
Captions for movies S1 to S5
References for SI reference citations

**Other supplementary materials for this manuscript include the following:**

Movies S1 to S5

### Supporting Information Text

#### Culturing *E. coli* for dead-end mother-machine measurements

After overnight growth, 2-4 ml of stationary phase cell culture was concentrated by centrifugation, in the presence of 0.5 μg/ml of bovine serum albumin (BSA), to a volume that was 3-10 times the volume of the pellet. Then 2-3 μl of resuspended culture was pipetted into the main flow channel of a dead-end microfluidic mother machine (see Design and fabrication of Dead-end mother machine microfluidic chips below). The cells were then allowed to populate the dead-end channels for 1-2 hrs. Once these channels were sufficiently populated, tubings were connected to the device, and a flow of fresh growth medium supplemented with BSA (0.5 μg/ml) was started. This medium contained no antibiotics. A flow was maintained at 5 μl/min throughout each experiment using an NE 1000 Syringe Pump (New Era Pump Systems, NY). To ensure a steady-state growth, the cells were left to grow in channels for at least 14 hrs (16 hrs for slow growth) before starting imaging.

#### Hyper/hypo osmotic treatment

For hyperosmotic treatments of E. coli, the medium was supplemented with different concentrations (0.1-1.0 M) of NaCl. For hypoosmotic treatments, DI water supplemented with 0.5 μg/ml of BSA was used. The osmolality value in each measurement was determined by a Vapro® 5520 vapor pressure osmometer (Wescor Inc, UT). Before applying hyper- or hypoosmotic shock, cells were imaged in regular media for 4 hours in mother machine devices. A hyper- or hypoosmotic shock was delivered to cells using a second syringe pump (NE-2000; New Era Pump Systems, NY), which was connected to microfluidic chips via a separate tubing. To prevent the high (or low) osmolality content reaching the cells prematurely, the tubing connecting to a syringe holding a high osmolality medium was backfilled with the regular medium before it was connected to the chip.

#### Rifampicin treatment

For rifampicin treatment, a separate tubing was connected to the microfluidic chip that contained M9 media with 300 μg/ml rifampicin. Care was taken for the drug from the tubing not to reach the cells before treatment by backfilling part of the tubing with the regular medium. First, cells were imaged in regular medium (4 hrs), then after a 60 s wait, the second syringe pump was started to flow the medium with rifampicin (5 μl/min). Cells were imaged in the presence of rifampicin for 20 min (25 min for slow growth). In experiments shown in Fig. 5, a hyperosmotic shock (0.2-1.8 Osm/kg) was delivered to cells after the rifampicin treatment. In these measurements, the hyperosmotic shock was delivered to the cells in the same way described in the previous section.

#### Squeezing measurements using microanvil devices

Microanvil is a pressure actuated microfluidic valve having a specific design (discussed in more details below). We made 112 microanvils on the same microfluidic chip. Of those, 56 were usable at the same time. Each microfluidic chip consisted of a glass coverslip, a PDMS (polydimethylsiloxane) elastomer layer of microfluidic flow channels, in which cells reside with growth media, and another PDMS elastomer layer of microfluidic control channels. The thickness of a flow layer was typically 10 μm, while the control layer was about 5 mm thick.

The flow channels and the control channels were arranged perpendicular to each other on the chip. There were 28 different flow channels on the chip. These channels all started from the same inlet and then spanned out in a binary-tree like structure. Each flow channel featured 40 patterns for microanvils. After the channels passed through the microanvil patterns they converged via binary-tree like structure to a single channel that leads to the outlet of the chip. Each chip contained two 50 μm wide and 21 μm high control channels. One of these channels was filled with 57% glycerol and connected to a pneumatic pressure line before the measurements started.

To squeeze a cell, pressure was applied to the control channel, causing the control channel to expand. This expansion caused the ceiling of the flow channel to be pushed toward the coverslip. The flow channels consisted of repeating patterns (microanvils) where the channel widened from either 2 or 3 μm to 10 μm. One of the repeating units is shown in Fig. 3A. In this wider region, the ceiling of flow channels featured a rectangular protrusion. The protrusions were 8 μm wide (perpendicular to flow channel) and 2 μm long (in the direction of the flow channel). Their height was 0.8 μm while the height of the flow channels was 2 μm. Away from the wider region, the flow channel gradually narrowed back to 2 or 3 μm width. Such a tapered shape helped in aligning cells relative to the flow channel axis.

At zero control pressure, the tip of the anvil was nominally 1.2 μm above the top of the coverslip. This was greater than the diameter of the cells, which ranged from 0.6 to 0.9 μm depending on growth conditions. The latter allowed cells to be loaded under the valve. To trap cells, we applied a low pressure (typically 22 psi) to the control lines while forcing a flow of cells and media through the flow channels. After cells were trapped, we stopped media flow and released control pressure. Squeezing measurements were started thereafter. The detailed fabrication procedure of microanvil devices is described below under SI Text **Fabrication of microanvil devices**.

### Design and fabrication of dead-end mother machine microfluidic chips

The fluidic circuitry in each chip consisted of a main channel for media supply and waste product removal and 600 dead-end channels connected to the main channel following a typical mother machine layout (Wang *et al.*, 2010). The main channels in all chips were 200 μm wide, and about 20 μm high. The chips that were used to image cells in different growth conditions had differently sized dead-end channels. For slow growth conditions, these channels had lengths of 22 μm, heights of 1.0 μm, and three different widths of 0.6, 0.7, and 0.8 μm. For moderately fast growth conditions, the dead-end channels had lengths of 17 μm, heights of 1.14 μm and two different widths of 0.8 and 0.9 μm.

The fabrication process for microfluidic devices was based on soft-lithography of polydimethylsiloxane (PDMS) elastomers. The process is described in further details in Yang *et al.* (Yang *et al.*, 2018).

### Fabrication of microanvil devices

Fabrication of microanvil devices was based on soft-lithography of multilayered monolithic PDMS elastomers (Unger *et al.*, 2000). Each layer of the microfluidic channels was molded using silicon (Si) wafer molds. The fabrication process for these molds is described in **Fabrication of Si molds** (below).

To make the control layer, a passivated wafer mold with the control layer pattern was placed underneath a homemade PTFE plate which had an opening of 3.1 inches (~80 mm) diameter in its center. This plate had a thickness of 1/4 inch (~6 mm). A layer of 10:1 (weight ratio of base:linker) mixture of PDMS (Sylgard 184 kit, by Dow Corning, MI) was poured on the wafer mold resulting in about 5 mm thick layer of PDMS. PDMS mixture in the holder was degassed in a desiccator at a pressure of about 0.2 atm. After degassing, the mixture was heated in a convection oven at 90°C for 15 min. The assembly was then removed from the oven and allowed to cool at room temperature for about 15 min. The Si mold was then separated from the PDMS and its holder. Access holes to control channels were punched without removing the partially cured elastomer from the PTFE holder.

For the flow layer, a degassed PDMS 10:1 mixture was spin coated on its wafer mold at 5000 rpm for 2 min. The coated wafer was then heated in a convection oven at 90°C for 12 min. Subsequently, the bonding surfaces of control and flow layers were activated using $O_2$ plasma treatment for 20 s. Then, the PTFE holder holding the control layer was connected to a micromanipulator. The control and flow layers were aligned relative to each other by using the micromanipulator while observing the layers in an optical microscope. After alignment, the layers were brought into contact using the vertical control of the manipulator. To strengthen the bond and to allow the PDMS to cure fully, the layers were further baked at 90°C for 15 min. The resulting monolithic bilayer was peeled from the flow layer mold. Then, individual chips were cut out and the access holes to flow channels were punched with a biopsy needle. The last step of assembly was to bond this structure to a #1.5 glass coverslip (Fisher Scientific). Prior to this bonding, glass and PDMS surfaces were activated using $O_2$ plasma treatment. The control channels in the assembled chips could withstand externally applied pressures of about 6 atm.

### Fabrication of Si molds

The fabrication of the Si molds for the mother machines follows the process described earlier (Yang *et al.*, 2018). Briefly, the patterns of dead-end channels were defined by a JEOL JBX-9300FS electron beam lithography system (JEOL, Japan) with ZEP520A, a positive tone e-beam resist (ZEON Chemical, Japan). After e-beam writing and resist development, a 15 nm chromium layer was deposited. Subsequently, the e-beam resist layer was lifted off using sonication in an acetone bath. A cryo-Si etch was carried out at −100 ℃ in an Oxford Plasmalab 100 inductively coupled plasma reactive ion etching system (Oxford Instruments, MA). The Cr layer acted as a mask for Si etching. The patterns for the larger flow channels were defined using photolithography of SU-8 2015 (MicroChem, MA). The reliefs that result from this step have a typical height of 20 μm.

Si mold of the flow layer of the microanvils was fabricated using three lithography steps. In the first step, the smallest flow channels and chip alignment marks were patterned by e-beam lithography. Then using an evaporated Cr mask, anisotropic smooth sidewall RIE etch of Si was carried out at 20°C. The result of this etching step was 2.0 μm high. For the second layer, a ~500 nm thick layer of ZEP520A was spun on the wafer at 2000 rpm for 60 s. With chip-by-chip alignment, the patterns of protrusions (microanvils) were defined on this resist layer by a second e-beam lithography step. ZEP520A was then developed, and the exposed Cr mask was removed by using sputter etching with Ar. Further etching of protrusions was carried out using the same anisotropic smooth sidewall Si RIE. Finally, a ~5 μm thick layer of a negative-tone photoresist was patterned for the wider portions of flow channels. These patterns were defined using photolithography.

Si mold for the control layer of the microanvils was defined using photolithography of SU-8 2015 (MicroChem, MA). The reliefs were about 21 μm thick.

All photomasks were written by a Heidelberg DWL66 laser lithography system (Heidelberg Instruments), with a 20 mm write head. The substrates of the photomasks were soda lime glass with a chrome layer covered by a photoresist.

All channel heights were measured from the Si mold. For the height measurements, a KLA-Tencor P-6 Stylus (KLA-Tencor Corporation, CA) profilometer was used. All wafer molds were silanized in a desiccator using (tridecafluoro-1,1,2,2-tetrahydrooctyl)-1-trichlorosilane (UCT Specialties, CA) for at least 15 min before using.

### Determination of crowder concentration based on the intensity of fluorescent reporters

The relative change in concentration of any species of crowders ($c/c_0$) was the same during osmotic shock and squeezing measurements. Therefore, one could use the relative change in concentration of cytosolic fluorescent label as a proxy for ($c/c_0$). The concentration of the label was proportional to the fluorescent intensity of the label from unit volume. As the cell shrinks, the cytoplasmic fluorescent reporter became more concentrated and resulting intensity increased in proportion to concentration assuming the cell width on average remains constant. Consistent with this assumption, our data showed that change in cytosolic width was less than 3% while length change was 40% at the highest osmotic shock measurement (Fig. 2C, D). Thus, $c/c_0 = I/I_0$, where $I$ was the average pixel intensity from the cell during the shock and $I_0$ before the shock. To determine this average intensity, first, the center line of the cell from one pole to another was determined manually. This centerline was then shortened by five pixels (0.55 µm) from both poles as these pole regions have different widths than the cell body. The centerline was then broadened by four pixels on each side along the short axes of the cell (total of 0.96 µm wide) and the sampled intensities from all these pixels were averaged, yielding an average intensity $I$. In case, the longitudinal profile was shorter than 12 pixels, only its middle 2 or 3 pixels were averaged and used in further analysis. To correct for bleaching, focus shifts, etc, $I/I_0$ ratios were further normalized by the ratio of total fluorescent intensities from the cell, $I_{\Sigma,0}/I_{\Sigma}$. Here $I_{\Sigma,0}$ is the total intensity before and $I_{\Sigma}$ during osmotic shock. These intensities were calculated using all pixels from the cell (broadened by six pixels on each side). This rectangular region covers a cell from pole to pole and was 1.4 µm wide. The final adjusted formula, which was used in calculations, was $c/c_0 = (I/I_0) \cdot (I_{\Sigma,0}/I_{\Sigma})$.

To analyze squeezed cells, the cell center lines were again first manually determined. These resulting curves were piece-wise linear. In determining *I*, the polyline was broadened to cover nine pixels (about 1 µm). The intensity line profile from this contour was then fitted to multiple exponential power functions to yield a smooth curve. Then, only values within the inflection points offset minus 5 away from corresponding centroids were taken into account for calculating mean cytoplasmic intensity *I*. Calculation of $I_{\Sigma}$ follows the same procedure as outlined for osmotic shock measurements.

### Coarse-grained Brownian Dynamics simulations

We modeled the *E. coli* chromosomes as a closed string of spherical beads. Several previous works have used similar approaches (Kim *et al.*, 2015, Shendruk *et al.*, 2015, Männik *et al.*, 2016). Each chromosomal bead had a diameter $a$. Estimations of this length scale are given below. Crowders were also represented by spherical beads. Their diameter was denoted by $a_c$. We carried out two series of simulations one having $a_c = 0.25a$ and the other $a_c = 0.4a$. The inner membrane of the cell was modelled as a cylinder with flat ends, as shown in Fig. 6A. The model defined two different constraints for this boundary. The chromosome experienced a boundary with a smaller radius and length than the boundary experienced by crowders. The inner boundary (for chromosome) had a length of $20a$ and a radius of $8a$. The difference in both diameters and lengths between the inner and outer cylindrical boundary was $2a_c$. These two boundaries were needed to reduce adsorption of chromosome beads to the inner wall. Alternatively, others have used single boundary but varied differently the magnitude of repulsive potential between the wall and the chromosomal beads and between the wall and the crowder beads to reduce absorption (Kim *et al.*, 2015).

Chromosome beads were linked to each other via a Finite Extension Nonlinear Elastic (FENE) potential:

$$U_{FENE}(r) = -\frac{1}{2}kR_0^2 \ln\left(1-\frac{r^2}{R_0^2}\right)$$

where $r$ was the distance between a given pair of nearest beads, $k$ was a positive constant, and $R_0$ was the maximum possible separation between that pair of consecutive beads. The excluded volume interactions between any pair of particles (one can be a chromosome bead, a crowder bead, or a constraint), (i, j) were modeled by a Weeks-Chandler-Andersen (WCA) potential, which was purely repulsive:

$$U_{WCA}(r) = \begin{cases} 4\varepsilon_0\left[\left(\frac{\sigma_i+\sigma_j}{2r}\right)^{12} - \left(\frac{\sigma_i+\sigma_j}{2r}\right)^{6} + \frac{1}{4}\right] & r < 2^{1/6}(\sigma_i + \sigma_j) \\ 0 & otherwise \end{cases};$$

where $r$ was the distance between particles $(i, j)$, $\varepsilon_0$ was a positive constant, and $\sigma_i$ was the diameter of particle $i$. All beads were set to have identical masses, $m$. During simulations, frictions and random forces were applied to every bead by a Langevin thermostat.

In earlier works effective diameter of chromosome beads, $a$ had been estimated (Kim *et al.*, 2015, Pelletier *et al.*, 2012, Männik *et al.*, 2016) to range between 50-400 nm. Following our previous work, we used here $a$ = 80 nm, which represents ~30 kb of dsDNA (Männik *et al.*, 2016). Therefore, 150 beads represented one genome equivalent of chromosome. The diameters of crowder beads correspond to $a_c = 0.4a = 32$ nm in one modelling series, $a_c = 0.25a = 20$ nm in the other. The inner cylindrical constraint corresponded length of 20a = 1.6 μm and diameter of 8a = 0.64 μm. These values approximately match the dimensions of a cell measured in slow growth conditions. Additional parameters used in the model were:

- Langevin temperature: $T = 1.0\ \varepsilon_0/k_B$;
- Langevin fraction: $\gamma = 0.1\ a^{-1}\sqrt{(\varepsilon_0/m)}$;
- WCA potential prefactor: $\varepsilon_0 = 4.11\times10^{-21}\ J$;
- FENE potential prefactor: $k = 30\ \varepsilon_0/a$;
- FENE maximum distance: $R_0 = 1.5a$;
- Difference between the diameters of inner and outer constraints: $2\ a_c$.
- Diameter used for constraints when calculating WCA potential: 0.

For each simulation, the chromosomal string of beads was initially placed in the vicinity of an inner constraint along a rectangular contour. The crowders were randomly placed in a cylindrical region inside that rectangle. To initialize the simulation, we set a cap to the net force experienced by each particle. This cap was gradually increased as the simulation progressed and eventually removed after 2.5 million simulation steps. After this initialization phase, the coordinates of all beads were recorded at every 50,000 simulation steps. In this way, we recorded all bead coordinates for a total of 600 frames. The results were derived from the last 500 of those 600 frames.

To determine the nucleoid lengths and widths from simulations, we projected the coordinates of chromosomal beads to a plane parallel to long axes of the cell. In such a way, the simulations were quantified comparably to the live cell images. After the projection of the beads to a plane parallel to long axes of the cell, we broaden each point by a Gaussian. The width of the Gaussian was chosen equal to the width of point spread function of our microscope ($\sigma$ = 130 nm). The latter was measured using CdSe quantum dots. We added the projections of one hundred consecutive frames from the model together and measured the nucleoid length and width from the resulted image in the same way we have been measuring the nucleoids in live cells (see *Image analysis* above). Hence, we used five measurements for both the length and the width of the nucleoid from each simulation. The mean values from these five measurements have been reported in Fig. 6.

All simulations were performed using ESPResSo package (Limbach *et al.*, 2006) and ran on the Advanced Computing Facility managed by the University of Tennessee, Knoxville.

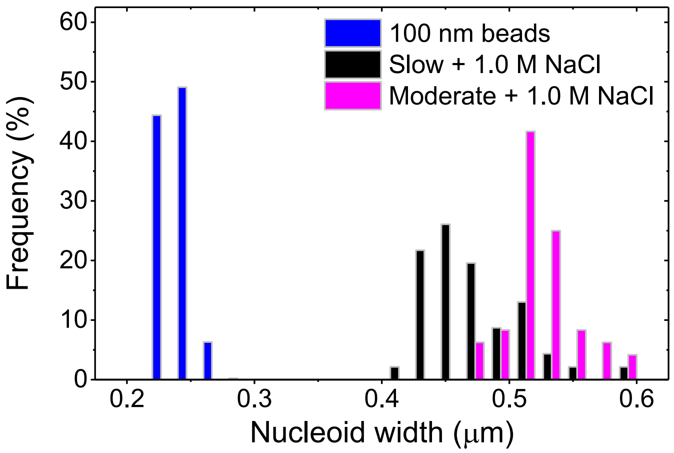


**Figure S1.** Comparison of nucleoid widths in slow (black bar, N = 46) and moderately fast (magenta bar, N = 43) growth conditions during 1 M NaCl shock to the widths of 100 nm fluorescent beads (blue bar, N = 491). 1M NaCl corresponds to an osmolality of about 2 Osm/kg (the highest concentration used in the measurements). The beads were from Life Technologies TetraSpeck Fluorescent Microspheres Sampler Kit. These beads were imaged with the same filter set and analyzed using the same analysis program as the nucleoids. The measured widths of beads correspond approximately to the width of the point spread function of the microscope. The latter width is about two times smaller than the width of the nucleoid in the most compressed conditions for the nucleoid.

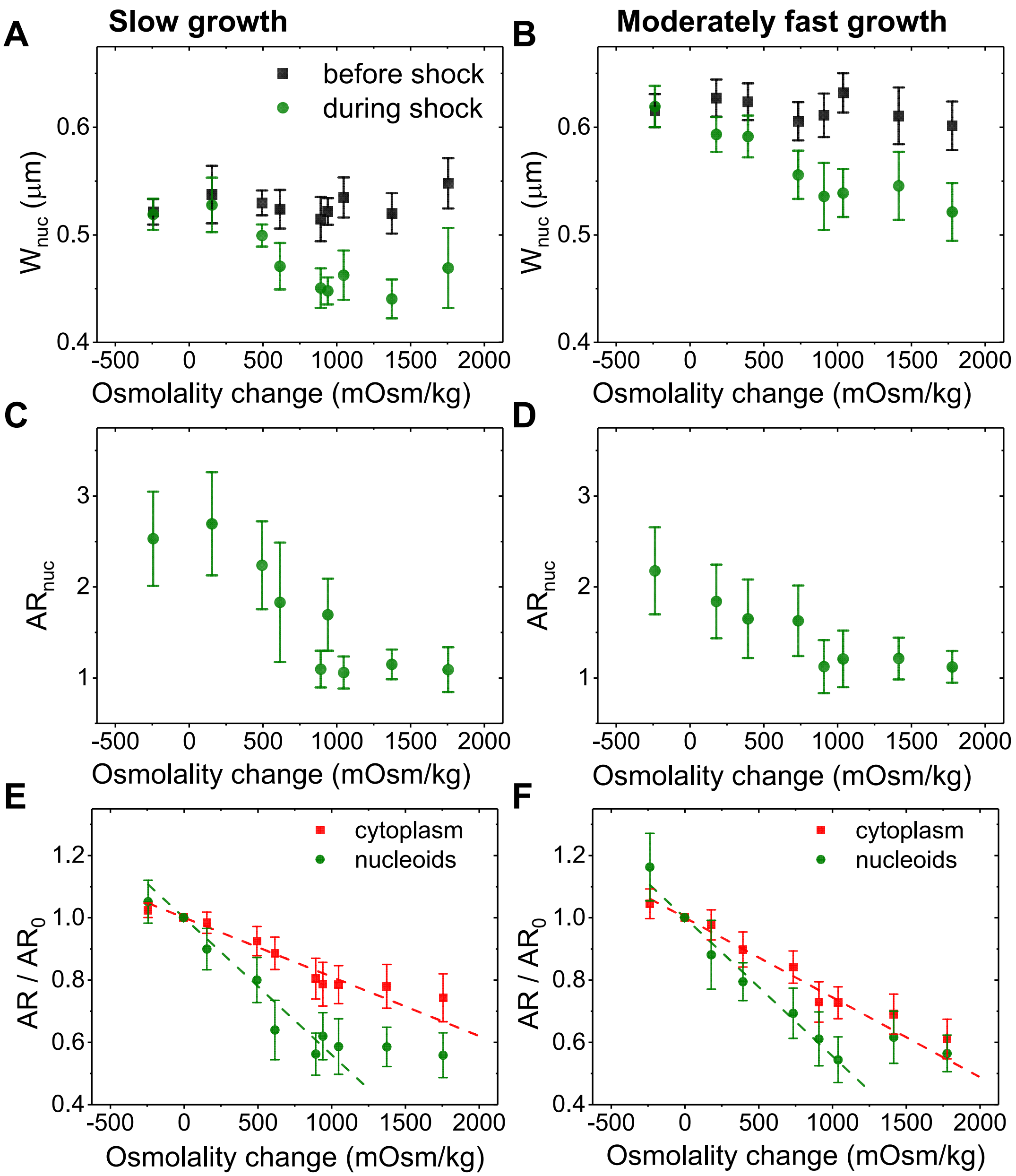


**Figure S2.** Measured aspect ratios of the nucleoids and nucleoid widths. (A) Nucleoid widths before (black) and about 1 min after an osmotic shock (red) in slow growth conditions. Shown are unscaled widths in µm. In Fig. 2, nucleoid widths are scaled by nucleoid widths before the shock. (B) The same in moderately fast growth conditions. (C) The aspect ratio of nucleoids during different osmotic shocks. The aspect ratio is defined as $\boldsymbol{L_{nuc}/W_{nuc}}$. (D) The same in moderately fast growth conditions. (E) Aspect ratios of nucleoids (green circles) and cytoplasm (red squares) right after osmotic shock relative to that prior the shock as a function of osmolality during the shock in slow growth conditions. (F) The same in moderately fast growth conditions. All error bars were std.

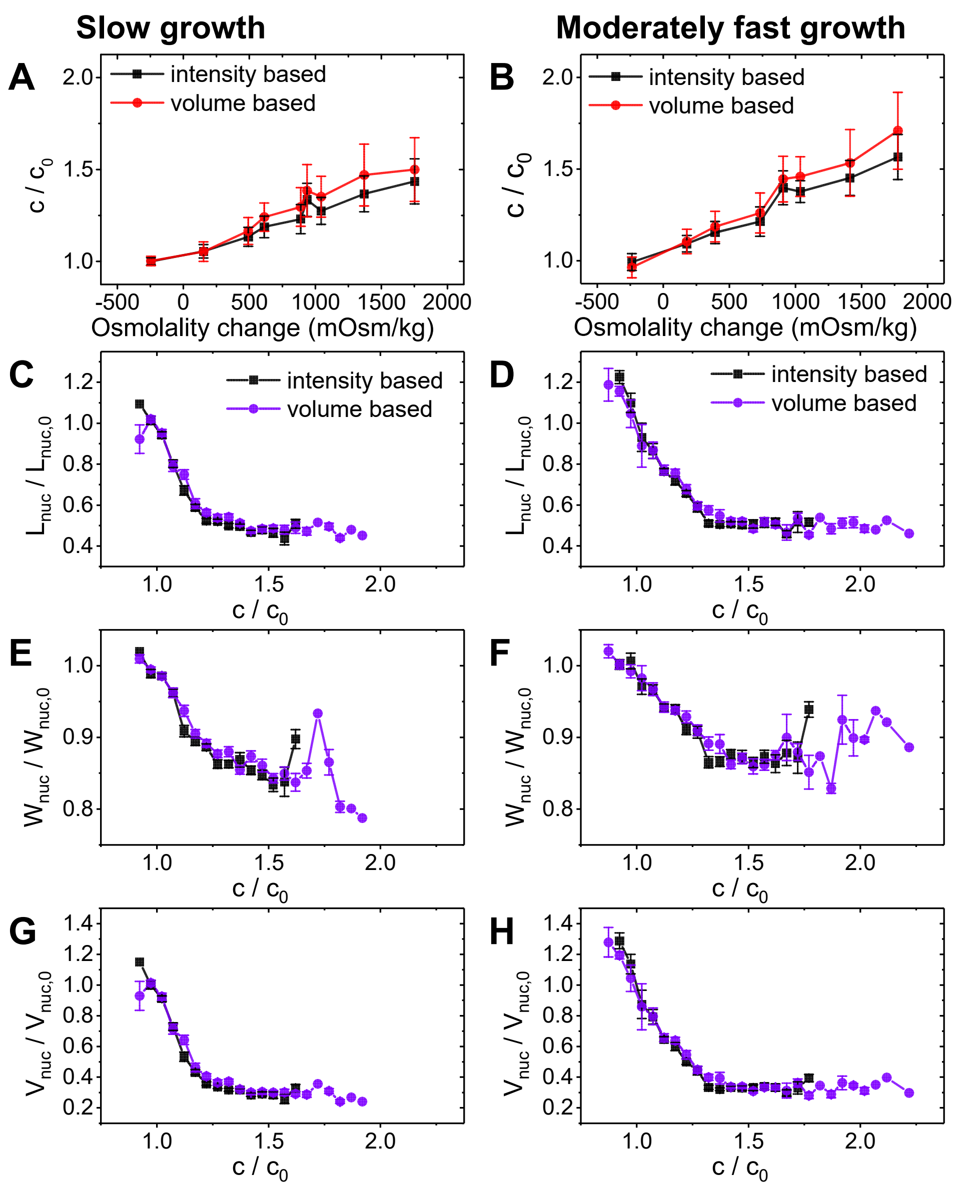


**Figure S3.** Comparison of cellular crowder concentrations determined using two different methods. The crowder concentration during osmotic shock ($c$) is determined relative to the concentration that cell maintains at regular growth medium ($c_0$). In intensity based method cytosolic tagRFP-T intensity is used as a proxy for the relative change of crowder concentration. The method is in further details described in SI Text: **Determination of crowder concentration based on intensity of fluorescent reporter**. In volume based method, the ratio $\boldsymbol{c/c_0}$ is taken equal to $\boldsymbol{V_{0,cyto}/V_{cyto}}$. Cytosolic volume, $\boldsymbol{V_{cyto}}$, is calculated based on the measured length and width of the cytosolic region assuming that this volume is a sphero-cylinder. (A, B) A relative crowder concentration as a function of external osmolality. (C, D) A relative change of the nucleoid length as a function of crowder concentration. (E, F) A relative change of the nucleoid width as a function of crowder concentration. (G, H) A relative change of the calculated nucleoid volume as a function of crowder concentration. Error bars are std in (A, B), and s.e.m. in (C-H).

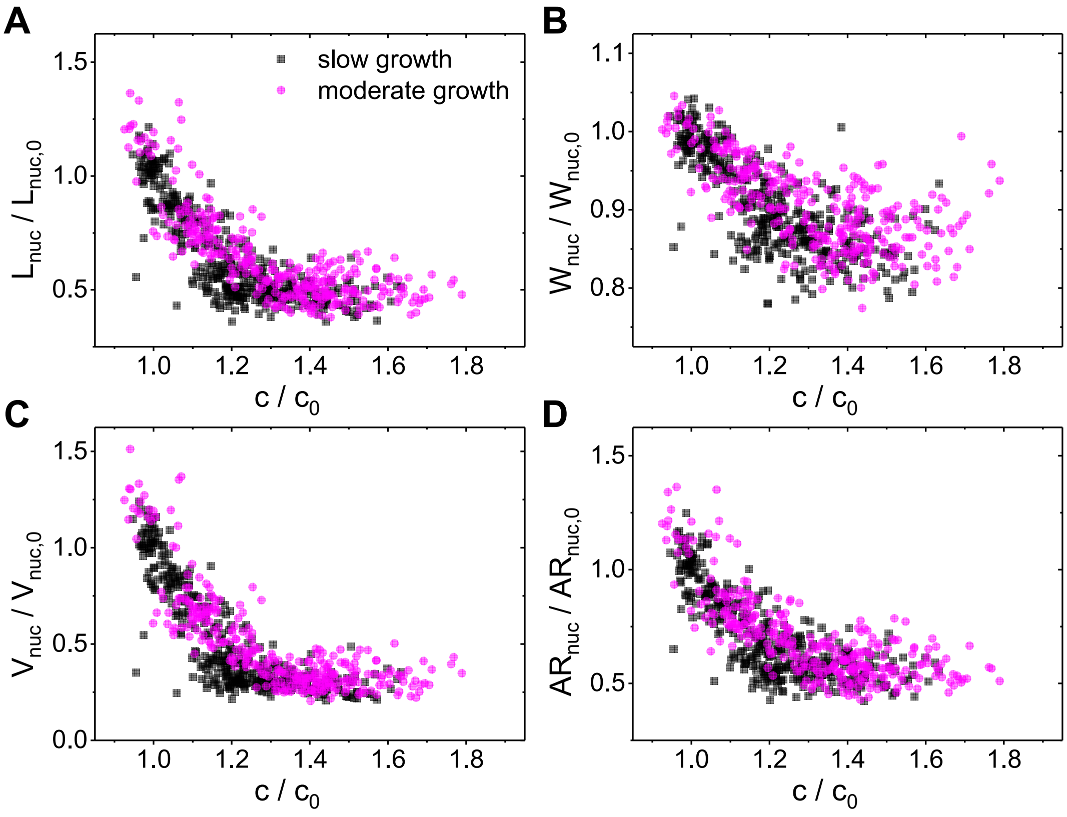


**Figure S4.** Raw data of normalized nucleoid lengths (A), widths (B), volumes (C) and aspect ratios (D) as a function of normalized crowder concentration in slow (black) and moderately fast (magenta) growth conditions.

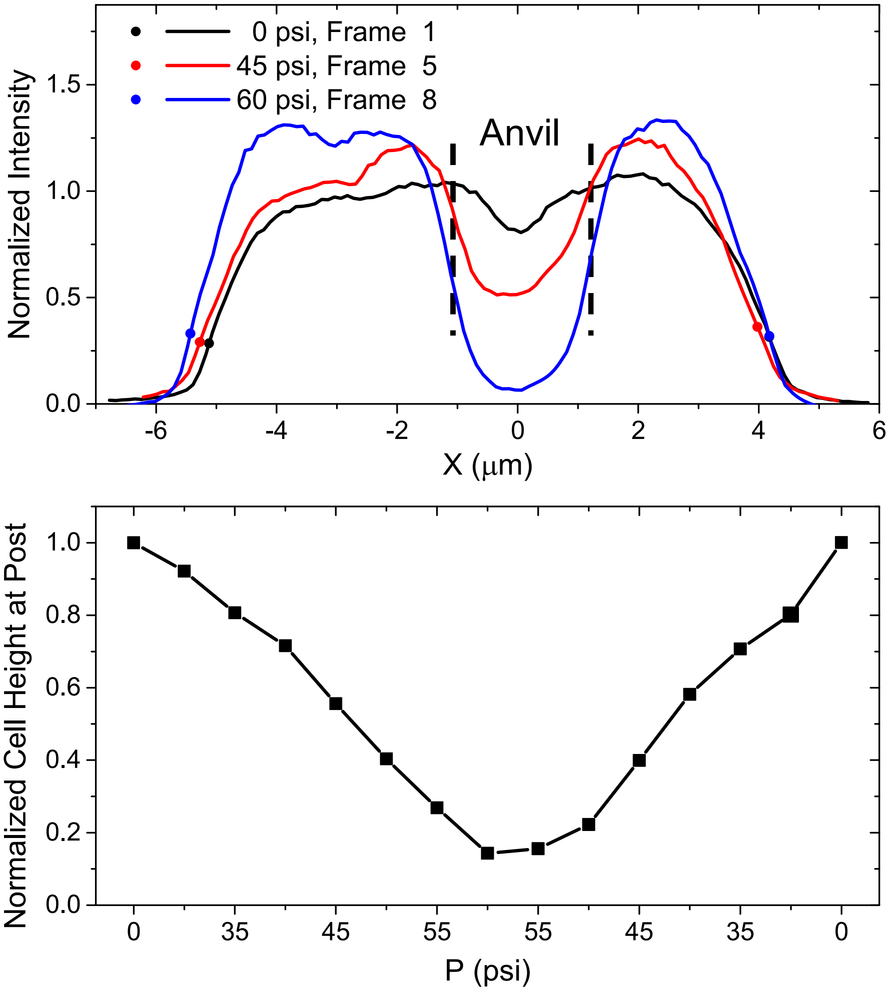


**Figure S5.** Top: Intensity traces of cytosolic mNeonGreen label along the cell length during squeezing measurements. Traces correspond to the cell shown in Fig. 3B. Bottom: Minimal intensity at the cell center at different pressures applied to the valve. All intensities are normalized by the intensity at the beginning of the measurements.

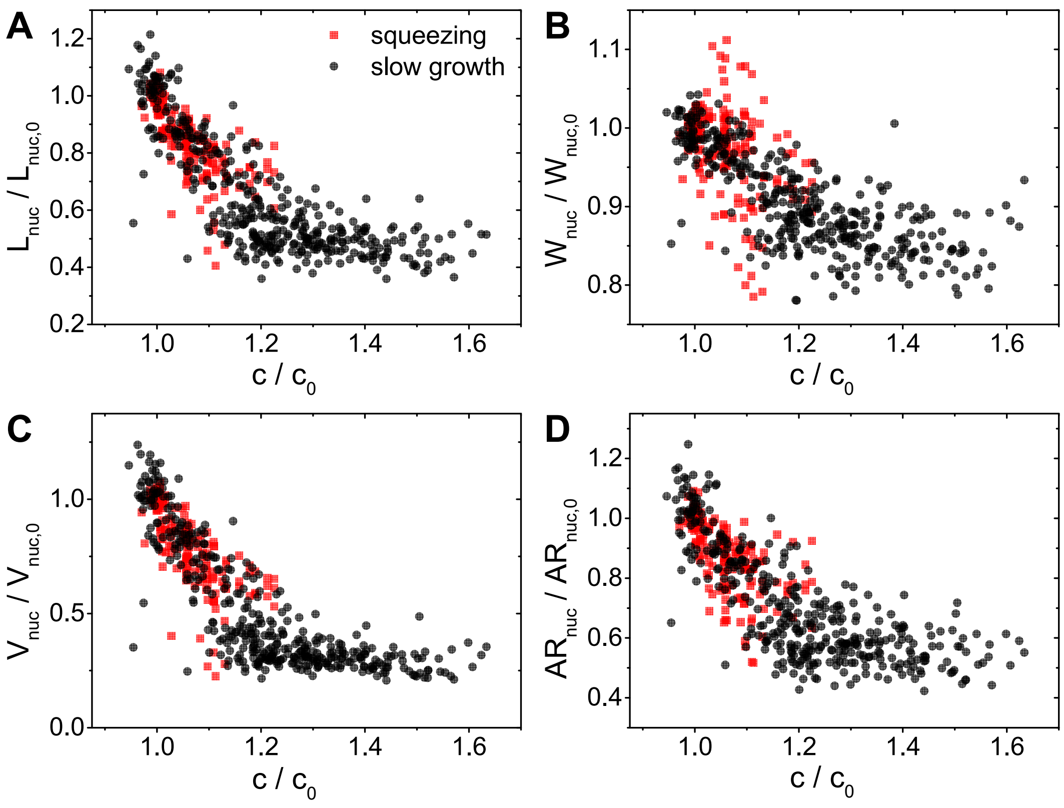


**Figure S6.** Comparison of raw data from squeezing measurements (red) to osmotic shock measurements (black). (A) Normalized nucleoid length, (B) width, (C) volume, and (D) aspect ratio as function normalized crowder concentration (A). Both squeezing and osmotic shock measurements have been performed in slow growth conditions.

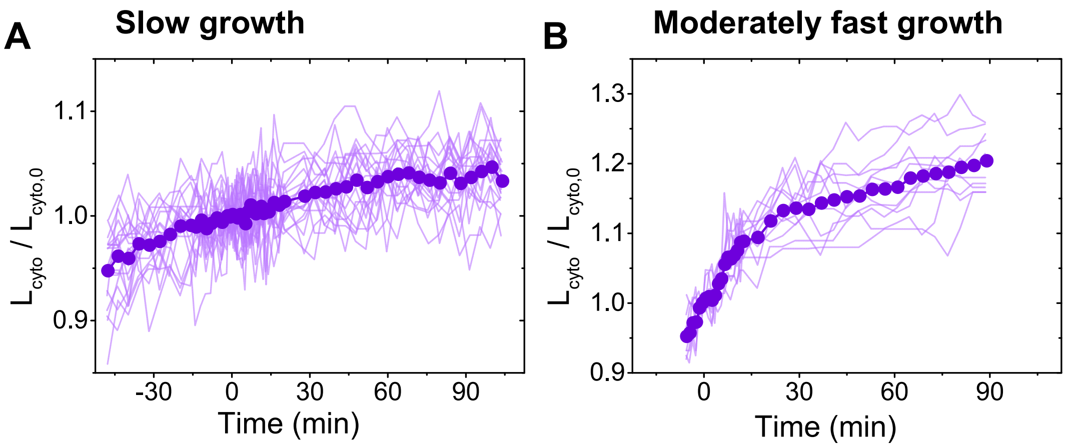


**Figure S7.** Growth curves prior to and during 300 µg/ml rifampicin treatment. Traces from individual cells are shown by thin light lines and population average curve by thick line with markers. Rifampicin circulation in microfluidic lines was started at 0 min. The drug is expected to reach cells within 1-3 min after the start of circulation. (A) Data for slow growth (N = 20), and (B) for moderately fast growth conditions (N=11).

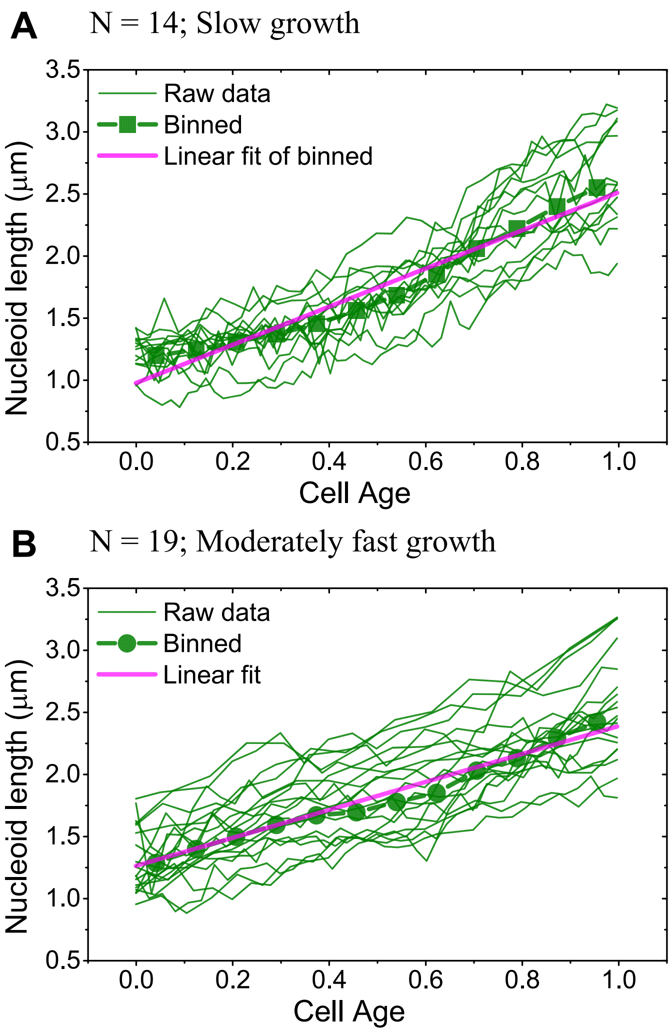


**Figure S8.** Nucleoid length as a function of normalized cell cycle time in slow (A) and in moderately fast (B) growth conditions. Thin curves are from individual cells. A thick solid line with filled circles is the population-averaged curve. A linear fit to the population is shown by the magenta line. In slow growth conditions, nucleoids elongate at the average rate of 6.8 nm/min and in moderately fast growth conditions at the rate of 12 nm/min. These values were used in extrapolating nucleoid lengths in osmotic shock measurements that followed rifampicin treatment (SI Fig. S9).

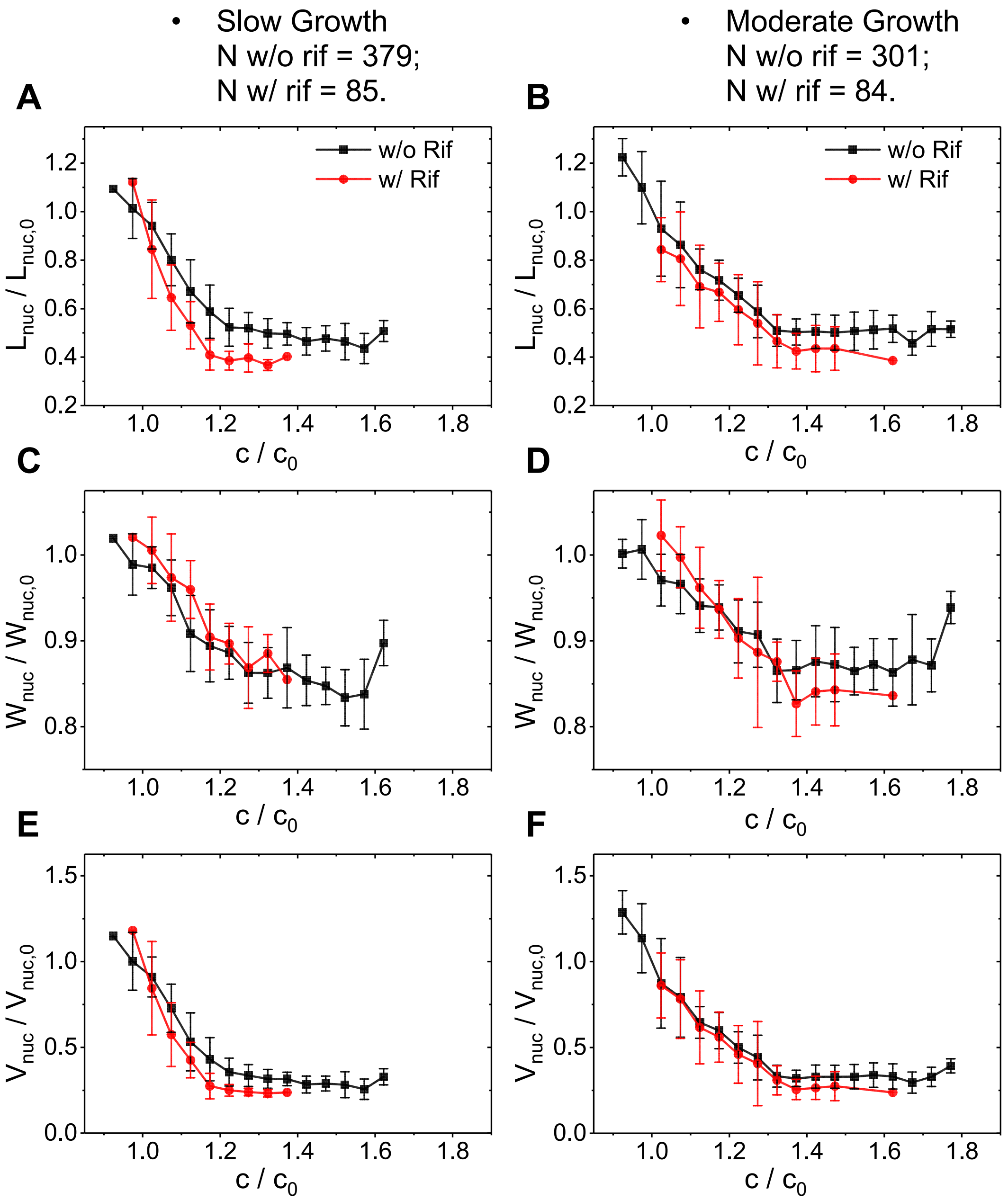


**Figure S9.** The effect of osmotic shock after rifampicin treatment calculated using extrapolated nucleoid lengths (based on Fig. S8). Nucleoid lengths, $L_{nuc}$, were extrapolated from the beginning of the rifampicin treatment (300 µg/ml) to the point 25 min later when the osmotic shock was administered. The extrapolated nucleoid size right after the osmotic shock relative to its size right before the shock is shown by red diamonds. For comparison, the compaction curve of the nucleoid during the osmotic shock but without rifampicin treatment is also shown (black squares; from Fig. 2G-F). (A, B) A relative nucleoid length as a function of relative crowder concentration. (C, D) The same for the relative nucleoid width, and (E,F) for the calculated volume.

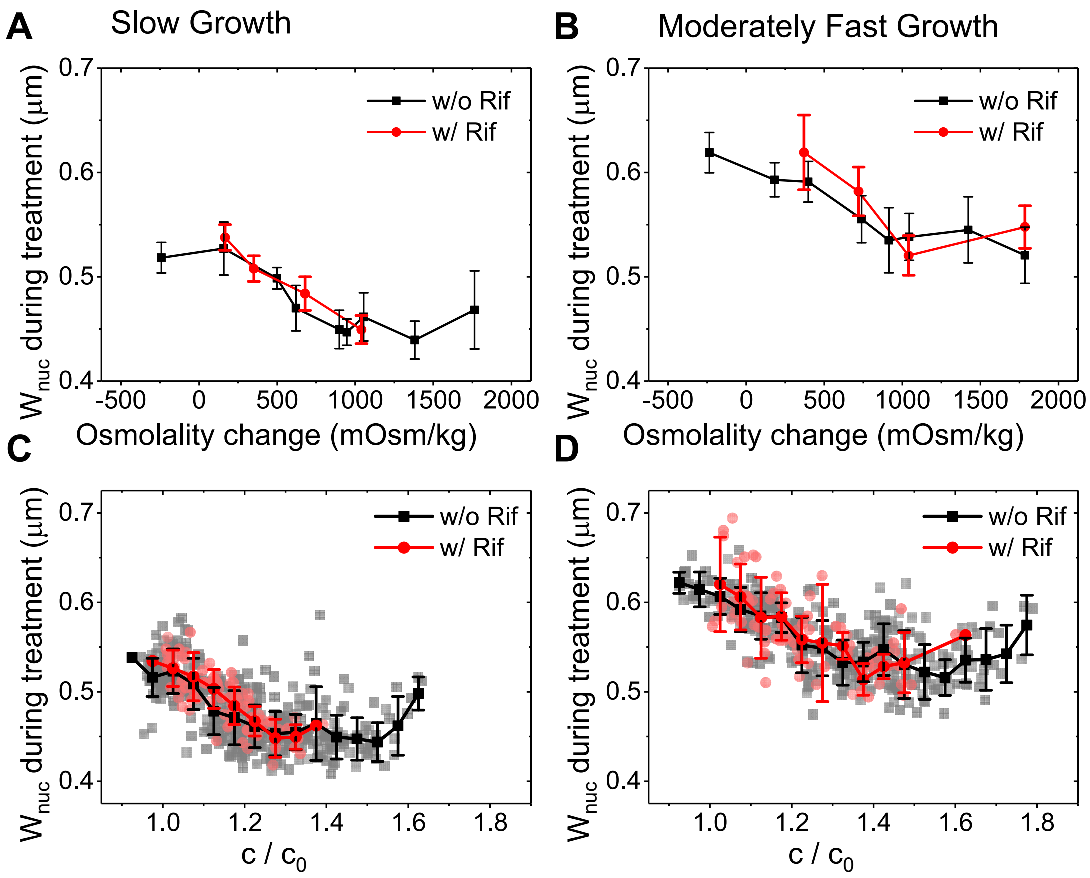


**Figure S10.** Nucleoid widths in μm before and during different treatments. (A, B) The average nucleoid width as a function of external osmolality in slow and in moderately fast growth conditions, respectively. Red circles correspond to 20 min 300 μg/ml rifampicin treatment of cells prior to osmotic shock and black squares to the same shock without rifampicin treatment. (C, D) The nucleoid widths as a function relative crowder concentration in slow and in moderately fast growth conditions, respectively. Overlaid to raw data are binned data points. Error bars for the binned data are std. In both slow and moderately fast growth conditions, the widths during osmotic shock are independent on whether a prior rifampicin treatment was carried out or not.

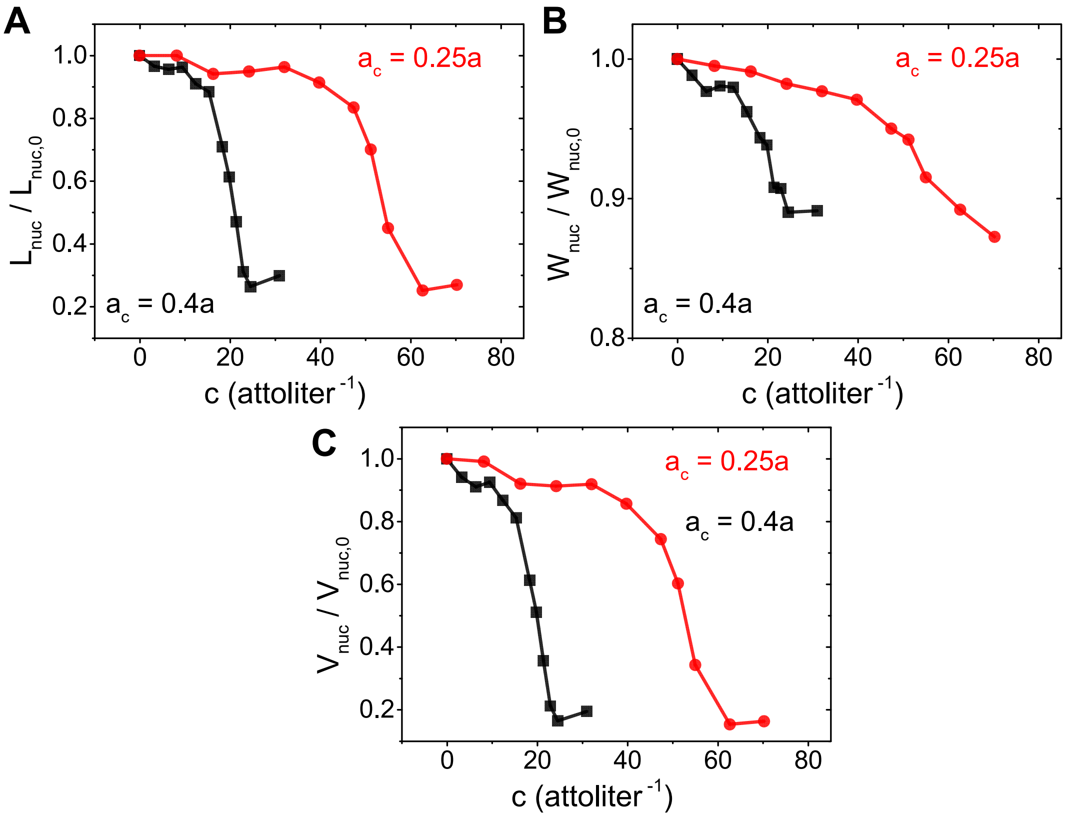


**Figure S11.** Nucleoid length, width, and volume from modeling as a function of crowder concentration. Curves for two different crowder sizes are shown. Concentration is calculated for a cylindrical cytosol that has a length of 1.6 μm (20*a*) and width 0.64 μm (8*a*). *a* = 80 nm.

**Table S1.** List of strains made and used in this study.

| Strain | Genotype | Used antibiotic concentration |
|---|---|---|
| JM57 | MG1655<br>*ΔleuB:: Pj231000 tagRFP-T::frt*<br>Δ*hupA::hupA-mNeonGreen-frt-kan-frt* | Plates/ liquid media: Kan 25 µg/ml<br>Microfluidics: no antibiotics |
| DY3 | MG1165<br>Δ*hupA::hupA-mCherry-frt*<br>*ΔleuB:: Pj231000 mNeonGreen- frt*<br>$P_{lac}$-*sulA cat* | Plates/ liquid media: CAM 25µg/ml<br>Microfluidics: no antibiotics |

**Table S2.** List of measurements. Columns from left to right: growth conditions, agent and concentration for osmotic and antibiotic treatment, osmolality prior the measurement, osmolality during the measurement and number of cells analyzed in each measurement (N).

| Regular Media | Treatment | Regular Media Osmolality (mOsm/kg) | Treatment Media Osmolality (mOsm/kg) | N |
|---|---|---|---|---|
| M9 + Gly | Water | 244 | 3 | 46 |
| M9 + Gly | 0.1 M NaCl | 239 | 397 | 45 |
| M9 + Gly | 0.3 M NaCl | 239 | 737 | 42 |
| M9 + Gly | 0.4 M NaCl | 239 | 860 | 46 |
| M9 + Gly | 0.5 M NaCl | 239 | 1137 | 47 |
| M9 + Gly | 0.55M NaCl | 239 | 1185 | 16 |
| M9 + Gly | 0.6 M NaCl | 241 | 1294 | 47 |
| M9 + Gly | 0.8 M NaCl | 252 | 1633 | 44 |
| M9 + Gly | 1.0 M NaCl | 252 | 2016 | 46 |
| M9 + Gly | Rifampicin | 241 | -- | 20 |
| M9 + Gly | Rif. + 0.1 M NaCl | 244 | 411 | 19 |
| M9 + Gly | Rif. + 0.2 M NaCl | 251 | 601 | 21 |
| M9 + Gly | Rif. + 0.4 M NaCl | 251 | 930 | 25 |
| M9 + Gly | Rif. + 0.6 M NaCl | 251 | 1290 | 20 |
| M9 + Glu + CAS | Water | 238 | 3 | 22 |
| M9 + Glu + CAS | 0.1 M NaCl | 240 | 423 | 23 |
| M9 + Glu + CAS | 0.2 M NaCl | 231 | 628 | 43 |
| M9 + Glu + CAS | 0.4 M NaCl | 242 | 981 | 43 |
| M9 + Glu + CAS | 0.5 M NaCl | 228 | 1142 | 36 |
| M9 + Glu + CAS | 0.6 M NaCl | 242 | 1286 | 42 |
| M9 + Glu + CAS | 0.8 M NaCl | 235 | 1657 | 44 |
| M9 + Glu + CAS | 1.0 M NaCl | 233 | 2019 | 48 |
| M9 + Glu + CAS | Rifampicin | 244 | -- | 11 |
| M9 + Glu + CAS | Rif. + 0.2 M NaCl | 239 | 609 | 21 |
| M9 + Glu + CAS | Rif. + 0.4 M NaCl | 245 | 964 | 22 |
| M9 + Glu + CAS | Rif. + 0.6 M NaCl | 245 | 1285 | 19 |
| M9 + Glu + CAS | Rif. + 1.0 M NaCl | 224 | 2009 | 22 |

**Table S3.** Slopes of linear fittings shown in Figure 2 (A-F).

| **Series of experiments** | **Measure** | **Fitted slopes** | | **Ratio of slopes** |
|---|---|---|---|---|
| | | Cytoplasm (kg/Osm) | Nucleoids (kg/Osm) | |
| M9 + Gly + NaCl | Length | -0.20 | -0.53 | 2.7 |
| | Width | -0.01 | -0.14 | 9.9 |
| | Volume | -0.24 | -0.72 | 3.0 |
| | Aspect Ratio | -0.19 | -0.44 | 2.3 |
| M9 + Glu + CAS + NaCl | Length | -0.27 | -0.52 | 1.9 |
| | Width | -0.02 | -0.13 | 6.1 |
| | Volume | -0.35 | -0.71 | 2.0 |
| | Aspect Ratio | -0.26 | -0.44 | 1.7 |

**Table S4.** Estimation of ribosome and protein numbers is slow and moderately fast growth conditions.

<table>
<tr><th></th><th>Slow growth</th><th>Moderate growth</th><th colspan="2">Explanation/Reference</th></tr>
<tr><td>Total number of ribosomes</td><td>15000</td><td>44000</td><td colspan="2">(Ehrenberg et al., 2013)</td></tr>
<tr><td>Polysome fraction</td><td>0.65</td><td>0.85</td><td colspan="2">(Mohapatra and Weisshaar, 2018)<br>(Dai et al., 2017)</td></tr>
<tr><td>Free 70S fraction</td><td>0</td><td>0</td><td colspan="2">(Mohapatra and Weisshaar, 2018),<br>(Dai et al., 2017)</td></tr>
<tr><td>Subunit fraction</td><td>0.34</td><td>0.15</td><td colspan="2">1-(Polysome fraction)</td></tr>
<tr><td>Ribosomes per polysome</td><td>10</td><td>13</td><td colspan="2">(Mondal et al., 2011)</td></tr>
<tr><td>Total proteins</td><td>2.1E+06</td><td>3.00E+06</td><td>Assuming the same concentration as in moderate growth</td><td>(Milo, 2013)</td></tr>
<tr><td>Cytosolic crowder fraction for proteins</td><td>0.40</td><td>0.20</td><td colspan="2">(Li et al., 2014)</td></tr>
<tr><td>Average Cell Volume [$\mu m^3$]</td><td>0.63</td><td>0.89</td><td colspan="2">Based on this work</td></tr>
<tr><td>Volume at Birth [$\mu m^3$]</td><td>0.43</td><td>0.61</td><td colspan="2">Based on this work</td></tr>
</table>

Cytosolic crowder fraction for proteins is based on Fig.5A in (Li *et al.*, 2014). Protein crowders belong to three sectors in this chart and include proteins involved in carbohydrate metabolism, nucleotide and amino acid metabolism, and in protein folding and decay. In fast growth conditions, these sectors account for 20% of all proteins. According to the text accompanying Fig.5A in (Li *et al.*, 2014), the contribution of these proteins can be expected to rise to 40% at the expense of decrease proteins involved in translation-related processes.

**Tables S5.** Estimated of crowder diameters ($a_c$), numbers (N), volume fractions ($\Phi_0$), concentrations ($c_0$) and crowding levels ($a_c^2c_0$) during normal growth conditions. % in the last columns shows crowding level for a given species relative to the total crowing level.

**Slow growth**

| | $a_c$ [nm] | N | $\Phi_0$ | $c_0$ [molecules/μm$^3$] | $a_c^2c_0$ [μm$^{-1}$] | % |
|---|---|---|---|---|---|---|
| Polysomes | 80 | 975 | 0.415 | 1.55E+03 | 9.90 | 0.20 |
| Subunits | 14 | 10200 | 0.023 | 1.62E+04 | 3.17 | 0.06 |
| 70S | 21 | 0 | 0.000 | 0.00E+00 | 0.00 | 0.00 |
| tRNA | 4 | 135000 | 0.007 | 2.14E+05 | 3.43 | 0.07 |
| Proteins | 5 | 8.49E+05 | 0.088 | 1.35E+06 | 33.71 | 0.67 |

**Moderately fast growth**

| | $a_c$ [nm] | N | $\Phi_0$ | $c_0$ [molecules/μm$^3$] | $a_c^2c_0$ [μm$^{-1}$] | % |
|---|---|---|---|---|---|---|
| Polysomes | 80 | 2942 | 0.886 | 3.31E+03 | 21.16 | 0.44 |
| Subunits | 14 | 13500 | 0.022 | 1.52E+04 | 2.97 | 0.06 |
| 70S | 21 | 0 | 0.000 | 0.00E+00 | 0.00 | 0.00 |
| tRNA | 4 | 405000 | 0.015 | 4.55E+05 | 7.28 | 0.15 |
| Proteins | 5 | 6.00E+05 | 0.044 | 6.74E+05 | 16.85 | 0.35 |

Estimation of crowder numbers

Following calculations are based on entries on Table S4

Number of Polysomes = (Total Ribosomes)*(Polysome Fraction)/(Ribosomes per Polysome)

Number of Subunits = 2*(Total Ribosomes)*(Subunit Fraction)

Number of tRNA = 9*(Total Ribosomes)  According to (Bremer and Dennis, 2008)

Number of proteins based on Table S4.

Estimation of polysome size

Polysome size is estimated from the diameter of 70S ribosome subunit (21 nm) and from the number ribosomes per polysome using Flory formula

$$a_{c,polysome} = a_{c,ribosome} N_{ribo/poly}^{3/5}$$

Number ribosomes per polysome, $N_{ribo/poly}$, values are listed in Table S4. This size reflects the effective excluded volume size of polysome in polysome-polysome interaction and exceeds significantly (60 times) the excluded molecular volume of the polysome components. It can be expected that this size is an overestimation. Clearly, Flory formula is solely an estimation rather than a numerically accurate value, especially in the limit of a small number of monomers. Moreover, crowding from proteins can be expected to compact polysomes. We choose here to overestimate the size of polysomes to be confident in our claim that polysomes do not have the predominant contribution to the compaction.

Estimation of subunit sizes

30S and 50S shell volumes have been estimated to be $1.3\times10^3$ $nm^3$ and $2.3\times10^3$ $nm^3$ (Bionumbers, N. R. Voss 'Geometric Studies of RNA and Ribosomes, and Ribosome Crystallization' 2006 p. 97). Although both complexes are oblate shape treating them as spheres yields effective radii 6.8 nm and 8.2 nm, respectively. Furthermore, we consider both subunits to be of the same size with an average diameter, 14 nm.

Estimation of tRNA sizes

The estimate is based on yeast Phe tRNA volume of 36 $nm^3$ (Bionumbers, N. R. Voss 'Geometric Studies of RNA and Ribosomes, and Ribosome Crystallization' 2006 p. 97). This yields an average diameter of 4 nm. The effective volume fraction is likely an underestimation taken the boomerang-like shape of the molecule.

Estimation of protein sizes

Clearly, proteins and protein complexes have a broad range of sizes--from the small ones, the diameters of ~2 nm, to GroEL chaperone complex, which has a molecular volume comparable to a 70S ribosome. The latter is estimated to be about 10 times less abundant than ribosomes though (Vendeville *et al.*, 2011). The available information remains too limited to calculate $c_i a_{c,i}^2$ for every protein. Therefore, we use 5 nm diameter as an average number as has been used in some estimations before (Odijk, 1998).

**Movie S1. Slow growing cells treated with 1.0 M NaCl.**

**Movie S2. Moderately fast growing cells treated with 1.0 M NaCl.**

**Movie S3. Slow growing cells squeezed using the microanvil device.**

**Movie S4. Slow growing cells treated with Rifampicin then 0.6 M NaCl.**

**Movie S5. Moderately fast growing cells treated with Rifampicin then 1.0 M NaCl.**
